# CcdA chaperones CcdB against irreversible misfolding and aggregation via a cotranslational folding mechanism

**DOI:** 10.1101/2024.10.15.618527

**Authors:** Priyanka Bajaj, Pehu Kohli, Raghavan Varadarajan

**Affiliations:** Molecular Biophysics Unit, Indian Institute of Science, Bangalore, India; Department of Bioengineering and Therapeutic Sciences, University of California San Francisco, San Francisco, CA, 94158, USA

**Keywords:** toxin-antitoxin, folding, stability, protein-protein interaction, chaperone, evolution

## Abstract

Cotranslational subunit assembly is thought to be a prominent feature throughout the proteome, but in bacteria, there are only a limited number of experimentally confirmed examples and most involve addition of extraneous tag sequences for experimental convenience. CcdA and CcdB are the antitoxin and toxin components, respectively, of the *ccdAB* operon. They assemble in a hetero-multimeric complex *in vivo*. Mutant phenotypes in a saturation mutagenesis library of CcdB were inferred from deep sequencing in two contexts, one when CcdB is expressed alone, and the other when CcdB is present in an operonic context downstream of its cognate antitoxin, CcdA. When expressed in the absence of CcdA, charged and polar mutations in the CcdB core cause the protein to misfold. However, many such deleterious mutations are rescued when expressed along with CcdA in the native operon. CcdA thus acts as a chaperone and relieves the folding defect in CcdBcotranslationally. Assembly efficiency and efficacy decreases when CcdA and CcdB are expressed from separate mRNAs relative to when they are expressed from the same polycistronic mRNA. Gene organisation in operons in bacteria may thus reflect a fundamental cotranslational mechanism that is important for effective assembly of protein complexes and can potentially buffer substantial genetic variation.

**Graphical Abstract:** 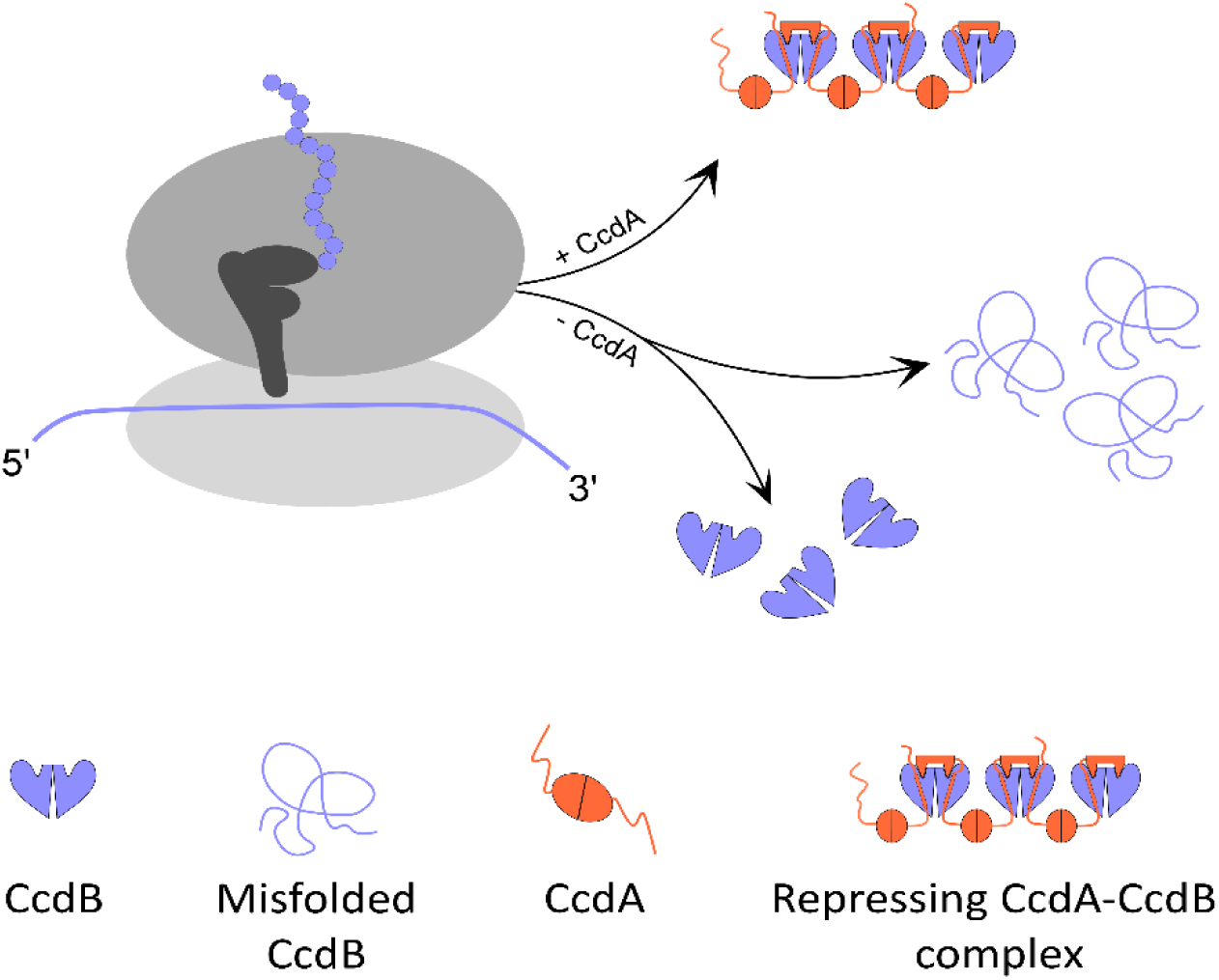

**In Brief:** Using the CcdAB Type II Toxin-Antitoxin system, the authors show that the antitoxin is able to suppress folding defects of the toxin, likely by processes involving cotranslational assembly.

**Highlights:**

- Deep mutational scanning is used to assess and compare the foldability of buried-site CcdB mutants in the absence and presence of CcdA.
- Folding defects of CcdB mutants are rescued in the presence of CcdA when expressed in an operonic context.
- CcdAB complex formation is enhanced when both the toxin and antitoxin components are synthesized from the same polycistronic mRNA, relative to synthesis from separate mRNAs.
- CcdA acts as chaperone both to assist correct folding, and to buffer mutations in CcdB, likely via cotranslational assembly mechanisms.

## INTRODUCTION

The faithful assembly of protein complexes in the cytosol is imperative to maintain cellular homeostasis (1). Oligomeric protein complexes were generally thought to be formed by diffusion and random collisions of subunits in the intracellular milieu (2), though there is increasing evidence that co-translational assembly occurs, especially for homodimeric proteins (3). Organization of bacterial genes into operons should promote assembly of heteromeric complexes as this results in increased local concentrations of both partners in the vicinity of the same mRNA molecule because of the presence of polysomes during protein translation. Theoretical analyses (4) provide support for this assertion, but there are limited experimental studies (5–7). Correct subunit assembly due to appropriate protein-protein interactions in the occluded and crowded cellular environment is important for protein stability and various cellular functions (8). Some of these biological functions include (i) enzymatic activities regulated by allosteric effects (9, 10), (ii) active-site formation by correct interaction of several catalytic/substrate-binding residues located in different subunits, (iii) generation of multifunctional enzymes by integrating functions compartmentalised in different enzyme subunits to form a single oligomeric enzyme (11) and, (iv) improving the stability of a monomeric protein upon oligomerization (12). Therefore, appropriate protein-protein interactions are crucial for correct subunit assembly, and folding intermediates are subjected to quality control checks by sequestration of proteases and chaperones (13). Improper folding may lead to formation of aggregates, which can result in several debilitating diseases (14–16). For instance, mutations causing misfolding of α-synuclein, tau protein or amyloid β-peptide can cause aggregopathies leading to severe diseases of the nervous system like Parkinson’s and Alzheimer’s (11, 15, 17). Hence, it is important to understand the molecular mechanisms involved in *in vivo* protein assembly, that discriminate correctly folded oligomers from aggregation prone monomeric units.

Protein assembly can occur either post-translationally or cotranslationally. In post-translational assembly, the protein assembly can either be near the site of synthesis known as cis-assembly or it can be at a distant site known as trans-assembly. Local confinement of assembly around translation sites promotes efficient and non-stochastic assembly (6). A growing body of evidence suggests that vectorial translation of proteins from the N to C terminus by ribosomes on a linear substrate affects the folding properties of single-domain, multidomain and oligomeric proteins (18). Many large proteins are not able to refold back or suffer from incorrect folding *in vitro* (19, 20) (21) (22). This problem is alleviated *in vivo* by establishment of a stable folding nucleus during cotranslational folding, which ensures that proteins fold into conformations that do not lead to aggregation or misfolding (23, 24).

The concept of cotranslational folding and assembly dates back to the 1960s, when researchers showed that nascent multimeric β-galactosidase enzyme possessed enzymatic activity before release from polysomes (5, 25). Several cytoskeletal elements including intermediate filament and sarcomere components are synthesised from localised mRNAs, suggesting that the synthesis of these proteins may occur at sites appropriate for function or assembly (26). There have also been several prior studies on the effects of synonymous mutations on the cotranslational folding properties of a protein (27–32) which suggest that codon usage can impact folding *in vivo*.

Cotranslational subunit assembly has also been studied, although how the folding defect of one of the interacting partners can be alleviated with the help of the other cognate partner is not well-understood. It is widely believed that mutations in the protein interior disrupt protein structure while mutations at exposed, active sites alter protein function (33–36). CcdA is the antitoxin and CcdB is the toxin component of the *ccd* operon (37). In isolation both proteins are homodimeric. Free CcdB exhibits its toxicity by binding to its cellular target, a type II topoisomerase, DNA Gyrase, thereby inhibiting DNA replication. CcdA neutralises the toxicity of CcdB by tightly binding to CcdB, inhibiting binding to DNA Gyrase (38). The *ccd* operon is also autoregulated. CcdA contains an N-terminal DNA binding and a C-terminal CcdB binding domain. When the CcdA:CcdB ratio is greater than 1, the two proteins form an extended (CcdA:CcdB)_n_ complex that binds to the operator region resulting in repression of the operon. However, under stress conditions Lon Protease degrades the labile antitoxin, and once the CcdA:CcdB ratio is less than one, this results in de-repression of the operon and fresh synthesis of *ccd* transcripts (39). The system thus enables the study of multiple protein:protein interactions in an operonic context *in vivo.* We have previously described construction and screening of deep mutational scanning (DMS) libraries of both the *ccdA* and *ccdB* components of the *ccd* operon (40, 41). In the present study, we further analysed the previously described *ccdB* DMS library (41) and carried out additional experiments that suggest that the CcdA:CcdB complexes assemble co-translationally, and that this provides an efficient mechanism for buffering otherwise deleterious mutations.

## RESULTS

### Rescue of misfolded, buried-site mutants in an operonic context

We have previously employed saturation mutagenesis coupled to deep sequencing to infer mutational phenotypes of CcdB in the absence of CcdA when expressed under the control of the pBAD promoter. Each CcdB mutant is assigned a variant score in the form of an MS_seq_ score (<u>M</u>utational <u>S</u>ensitivity score obtained from deep sequencing) that ranges from 2 to 9, which is a measure of its Gyrase binding activity (Figure S1 A). A mutant with a high MS_seq_ score implies that cells transformed with the corresponding toxin mutant survive upto high inducer (arabinose) concentrations. This is likely because the mutation results in decreased amounts of active CcdB, and only when the protein is highly overexpressed is there a sufficient amount of active CcdB to kill the cell (42). Thus, *in vivo* activity decreases going from an MS_seq_ score of 2-9 (43). The same site-saturation mutagenesis library was constructed in its native operon and was first transformed in the CcdB sensitive strain. Each mutant was assigned a variant score in the form of RF^CcdB^ (<u>R</u>elative <u>F</u>itness^CcdB^) which is a measure of free CcdB toxin levels in the cell relative to WT, higher levels of free CcdB competent to bind and poison DNA Gyrase, lead to enhanced cell death and low values of RF^CcdB^ (41). In contrast, a high RF^CcdB^ score is associated with lower levels of free CcdB and enhanced cell growth of cells transformed with the mutant operon relative to WT (Figure S1 B).

The site-saturation mutagenesis library of CcdB generated in its native operon in the resistant strain was also transformed in the RelE reporter strain to probe the ability of CcdB mutants to bind CcdA (Figure S1 C). The RelE reporter strain harbours a *relE* gene downstream of the *ccd* promoter, consisting of consensus Shine-Dalgarno (SD) sequence in pBT vector along with the *ccd* operon in pUC57 vector. Each variant was assigned a variant score in the form of RF^RelE^ (<u>R</u>elative <u>F</u>itness^RelE^). RF^RelE^ is a measure of the efficiency of CcdA-CcdB complex formation.

A high RF^RelE^ score implies that CcdB forms a tight complex with CcdA, that is able to bind the ccd operator and repress transcription (41). We looked at the frequency distribution of RF^RelE^ scores for the entire dataset, specifically for buried-site mutants and for the stop codon mutants, i.e., a construct with a non-functional toxin (Figure S2 A). The median of the distribution for the entire dataset lies close to 1 (41), an expected observation as a large fraction of the mutants in the other screen (-CcdA) gave an MS_seq_ score of 2 (44), suggesting that the majority of the mutants in both the screens display a phenotype similar to WT. We observed that RF^RelE^ scores for all the stop codon mutants were highly variable. Careful examination of the individual stop codons revealed that mutations to TAA were mainly concentrated at the derepressing end (RF^RelE^<1) while the TAG stop codons spanned the entire range of RF^RelE^ scores (Figure S3). Since TAA is the most commonly found stop codon in prokaryotic genomes (45), it likely terminates translation efficiently, resulting in non-functional CcdB protein fragments. The reason for the wide range in RF^RelE^ scores for TAG stop codons is currently unclear. It is possible that for TAG stop codons, either the termination codon is read-through (46) or the presence of premature termination codons might result in ssr tagging of proteins and possibly degradation of RNA (47, 48). Therefore, we analysed only nonsense mutations to TAA, considering it as a negative internal control, for which the median of its distribution is 0.52, i.e., showing a derepressing phenotype (Figure S2 A). Interestingly, despite the fact that in a non-operonic context, many mutations to buried residues are poorly tolerated and result in loss of activity because of a decrease in the amount of correctly folded protein (43, 44), here in the operonic context, 57% (308/541) of the buried-site mutants display a phenotype similar to WT, illustrated by the median of the distribution of the RF^RelE^ scores of buried-site mutants being close to 1.

In our previous study (41), we were able to successfully differentiate between non-interacting and interacting residues crucial for CcdA and DNA Gyrase binding, as well as identify buried and exposed residues within the protein structure based on RF^RelE^ and RF^CcdB^ score distributions (Figure S2)(41). This approach provided valuable insights into the diverse roles of residues in protein-protein interactions and their impact on overall protein function.

### Comparison of CcdB activity of buried-site mutants in the absence and presence of CcdA

We next compared phenotypes of individual buried site mutants when expressed in the presence and absence of CcdA. Mutations to charged, polar or glycine residues in the core of the protein show high MS_seq_ values ranging from 6 to 9 (Figure S4 A), implying that these buried-site mutants, when expressed in the absence of CcdA are impaired in binding to Gyrase, likely due to misfolding. We have previously shown that most buried site mutants with high MS_seq_ values are targeted to inclusion bodies when CcdB is expressed in the absence of CcdA (44). Burial of charged and polar groups in the non-polar interior of the protein destabilises the protein, and the degree of destabilisation depends on the relative polarity of the group. Mutations to glycine likely increase flexibility, and large to small substitutions result in an interior destabilising cavity in the core (44, 49, 50). Surprisingly, the majority of these mutants when expressed together with CcdA in an operon show RF^RelE^ scores similar to WT and, in some cases, such as for T65G, A93K, it is as much as two-fold higher than the WT (Figure S4 B). Even highly destabilising mutations at buried sites such as hydrophobic to charged substitutions are apparently rescued in the operonic context (Figure S4 B). Many buried site mutants are neutralised by CcdA as shown by high RF^CcdB^ scores (Figure S4 C). However, 19% (72/382) of buried-site mutants in their operonic context still exhibited significantly high levels of RelE toxicity (RF^RelE^<0.7) (Figure S2 C). In several cases, mutations to aspartate likely remain misfolded even when they are present in an operonic context (Figure S4 A and B).

We have previously characterised the thermal stability of 34 buried CcdB mutants (44). We calculated the difference in thermal stability of the mutant with respect to WT (ΔT_m_) and compared it with the corresponding MS_seq_, RF^RelE^ and RF^CcdB^ scores (Figure S4 D). Consistent with our previous analysis, many destabilised buried-site mutants that display a strongly inactive phenotype in the screen where CcdA is absent (4<MS_seq_<8), show a phenotype similar to WT in the screen in which CcdA is present in the operonic context, delineated by RF^RelE^ scores greater than 1. However, some mutations (L36K, L36A and V54D) show high MS_seq_ (4–8) and RF^CcdB^ (4–7) scores, and low RF^RelE^ (≤0.7) scores, indicating that these mutants remain misfolded even in their operonic context, because of which they lose binding to both CcdA as well as Gyrase (Figure S4). These analyses suggest that in several cases, the folding defect of CcdB is rescued in the presence of its interacting partner, CcdA, when both are present in the context of the native operon.

### Comparative solubilities of CcdB mutants when expressed alone and in an operon validate phenotypes seen for buried-site mutants

Expression from the *ccd* promoter results in very low levels of protein. Hence, we made a few constructs of buried-site *ccdB* mutants placed downstream of *WT ccdA* in the arabinose inducible pBAD vector. In these constructs, *ccdA* and *ccdB* coding sequences are expressed in an operonic context but with the native *ccd* promoter replaced by the stronger arabinose inducible P_BAD_ promoter. In this construct, there is low but still clearly detectable expression for CcdB. However, CcdA expression is too low to detect reliably in the operonic context. To understand how the mutants behave in comparison to when they are present without CcdA, we also constructed the same CcdB buried-site mutants alone in pBAD24 vector. Details of mutant phenotypes (MS_seq,_ RF^RelE^ and RF^CcdB^) and residue burial (%Acc and Depth) are mentioned in Table S1. We carried out *in vivo* solubility assays to examine the effect of CcdA expression on suppression of folding defects in CcdB mutants. For buried-site mutations with high MS_seq_ scores (4–9), in the absence of CcdA, the major fraction of the induced protein aggregated to form inclusion bodies, thereby going into the pellet fraction. However, for the same mutants in the presence of CcdA that gave RF^RelE^∼1 or RF^RelE^>1, a large fraction of the induced protein was observed in the supernatant, suggesting that in the presence of CcdA, the folding defect of many highly destabilised CcdB mutants is relieved (Figure 1 A, B and S5). V18R, L36R, and L36D mutants have high MS_seq_ scores. As expected, L36R and L36D exhibit low RF^RelE^ values, while V18R shows a paradoxically high RF^RelE^ score despite appearing degraded in the operonic context. In the absence of CcdA, a significant fraction of all three mutants is found in the pellet. However, they do not show any detectable bands in the operonic context, likely due to protein degradation (44). *In vivo* solubility estimates for 12 buried-site mutants both in the absence and presence of CcdA, in the majority of cases, validate our deep sequencing results (Figure 1 B and C). Given the low expression seen in the operonic constructs and the faint bands observed in Coomassie-stained gels, these results were subsequently confirmed through Western blotting experiments as described below.

**Figure 1:**
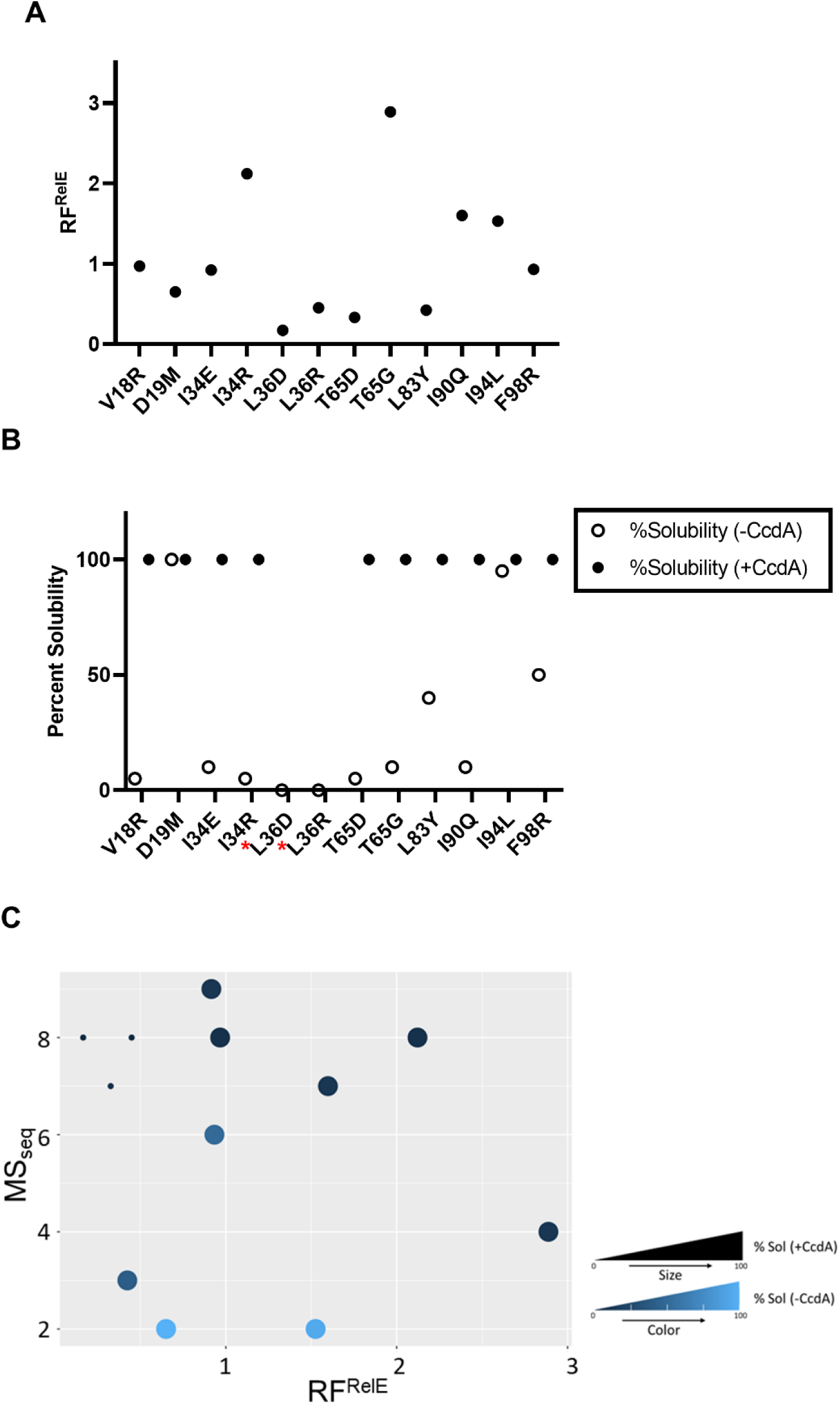
Expression and *in vivo* solubility of selected buried-site CcdB mutants in the absence and presence of CcdA, validate phenotypic effects inferred from deep sequencing data. (A) RF^RelE^ (+CcdA) obtained from deep sequencing for selected mutants. (B) The percent solubility which is an estimate of the fraction of the induced protein found in the supernatant both in the absence of CcdA (-CcdA) and in the presence of CcdA (+CcdA). (* indicates no detectable band in the operonic context) (C) The values of the percent solubility which is an estimate of the fraction of the induced protein found in the supernatant, both in the absence of CcdA (-CcdA) and in the presence of CcdA (+CcdA) as well as the corresponding MS_seq_ and RF^RelE^ values for all mutants are summarised in a scatter plot. The size and the color of each dot represent values of % solubility in the absence and presence of CcdA, respectively, for a given mutant.

### Expression of CcdB in operonic and non-operonic contexts

To understand whether the rescue of CcdB by CcdA is a cotranslational or a post-translational event, we constructed WT CcdB and CcdB mutants along with WT CcdA under identical but separate IPTG inducible T7 promoters separated by a strong synthetic tZ terminator (51) in the pET-Duet-1 vector (Figure S6 A). In this vector, CcdB and CcdA will therefore be synthesized from separate mRNAs. ccdA and ccdB were also cloned in an operonic context where the ccdB was fused at its C-terminus to a His tag (Figure S6 B). To compare the expression of CcdB WT in an operonic versus non-operonic contexts, SDS-PAGE and Western blot analyses were performed (Figure 2 A, and B and S7). Western blot results revealed a significant increase in the fraction of WT CcdB in the pellet, from 0.11 to 0.62 in the non-operonic relative to the operonic context.. For the T65G and T65D CcdB mutants, the fraction in the pellet increases from 0.08 and 0.08 in an operonic context to 0.62 and 1, respectively, in a non-operonic context, similar to the pattern observed with CcdB WT. The I34E CcdB protein is not detectable in the pellet in operonic context but, when expressed in a non-operonic context, shows clear targeting to inclusion bodies, resulting in its presence in the pellet. For the I90Q mutant, the fraction in the pellet increases drastically from 0 in an operonic context to 0.8 in a non-operonic context (Figure 2 C). The induction of the V18R mutant of the CcdB protein in the non-operonic context is minimal, and it remains misfolded, being entirely in the pellet fraction (Figure 2B, C). In the operonic context, no bands were found in either the supernatant or the pellet fraction upon Western blot analysis (Figure 2A). The T7 promoter used in the non-operonic context is stronger than the pBAD promoter used in the operonic context. Thus, the former will lead to higher expression of the V18R mutant. When CcdB and CcdA are expressed from different RNAs, the efficacy and the efficiency of the chaperonic effect of CcdA on CcdB diminishes, pointing to the involvement of cotranslational assembly in formation of the CcdAB complex., The chaperonic effect of CcdA in the operonic context was also examined in the context of Lon protease activation by thermal stress. Here, while WT CcdB and T65G maintained similar solubility levels at both 37 and 42°C, I90Q and I34E mutants (Figure S8 showed comparable amounts in both soluble and insoluble fractions at 42°C, unlike at 37°C where they were primarily in the supernatant (Figure 2A). These findings highlight the impact of protease activation on the CcdA-CcdB interaction.

**Figure 2:**
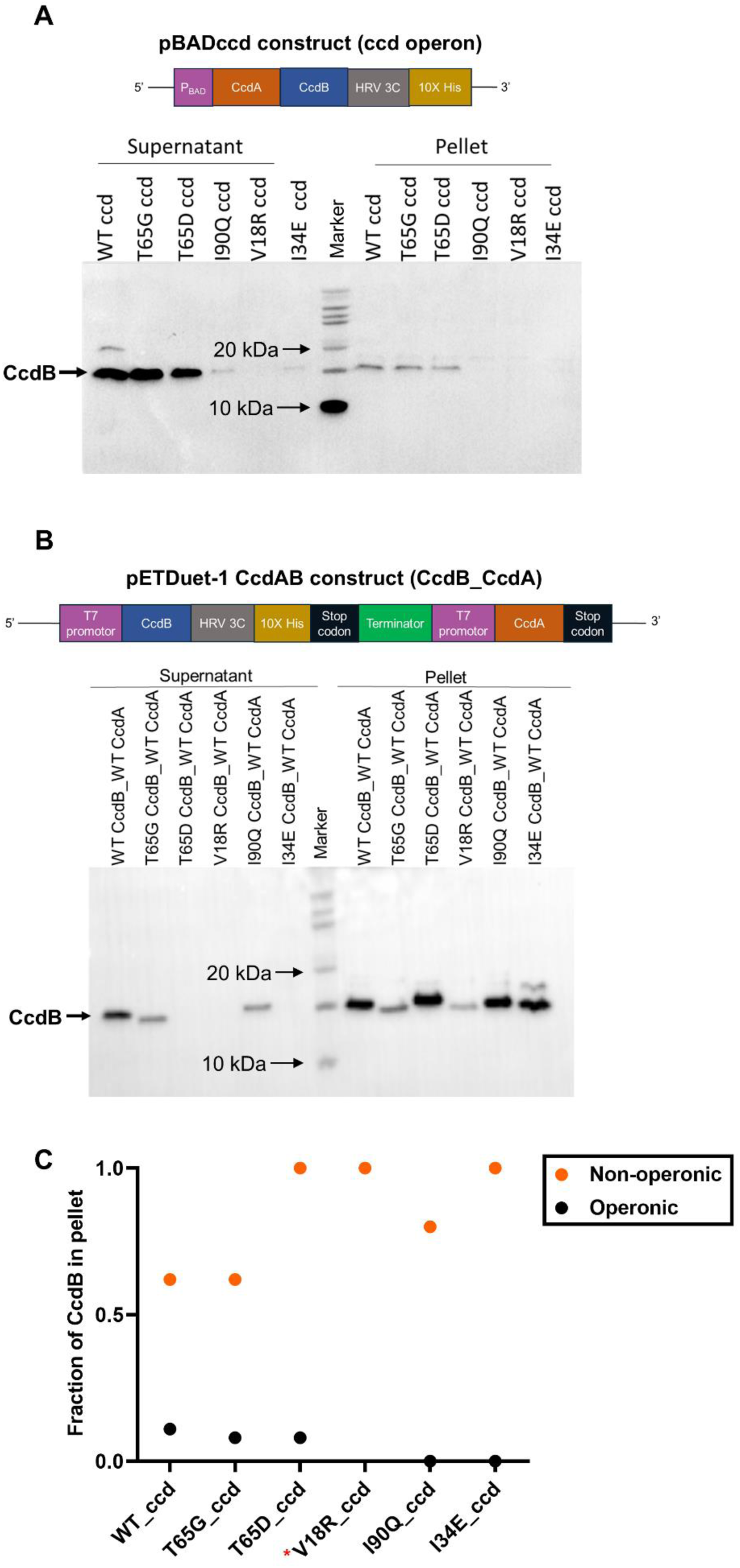
Comparison of CcdB protein expression of WT and destabilized mutants when CcdB and CcdA are expressed from the same and different mRNAs, respectively. CcdB and CcdA are expressed from (A) the same promoter and (B) separate but identical promoters separated by a strong terminator. The CcdB protein band is indicated by a black arrow in the western blot detecting His-tagged CcdB. The folding defect is not rescued when CcdB and CcdA are expressed on separated promoters as most of the CcdB is seen in the pellet fraction, suggesting CcdA chaperone’s activity of CcdB is likely via cotranslational folding processes. (C) Comparison of fraction of CcdB in pellet (amount of CcdB in pellet/total mount of CcdB) in operonic and non-operonic contexts. (* represents no detectable band in the operonic context)

### Estimation of protein yield of CcdB buried-site mutants when expressed in an operonic context

The CcdB mutants cloned with a C-terminal His-tag present in the context of the *ccdAB* operon were purified using Ni-NTA affinity chromatography (Figure S6 B). The fold change in protein yield of CcdB mutants relative to WT CcdB was estimated when CcdA and CcdB are expressed in the operonic context from pBAD24 vector. CcdB and CcdA bands with molecular weight of 14.25 kDa and 8.37 kDa, respectively, are visible (Figure 3 A). We quantitated the protein yield of the purified mutants relative to WT by subjecting them to SDS-PAGE and subsequently probing with anti-His antibody to detect the CcdB protein band on a Western blot (Figure 3 C). CcdB T65G (RF^RelE^ = 2.89) and CcdB I90Q (RF^RelE^ = 1.6) have almost 1.5 times the yield of CcdB WT (RF^RelE^ = 1). While CcdB T65D (RF^RelE^ = 0.33) and CcdB I34E (RF^RelE^ = 0.92) have a protein yield half that of CcdB WT (Figure 3 D). Surprisingly, for V18Rccd (RF^RelE^ = 0.97) neither CcdB nor CcdA protein band was detectable. Some extra bands other than the desired protein bands were also observed (Figure 3 A). These could either be chaperones that assist the cotranslational folding processes, or proteins that bind non-specifically to the Ni-NTA column. The ratio of copurified CcdA with CcdB was determined, and values ranged from 0.36 to 0.16 for the different mutants (Figure 3 B). The identity of the purified protein complexes was further confirmed by ESI Mass Spectrometry (Figure S9).

**Figure 3:**
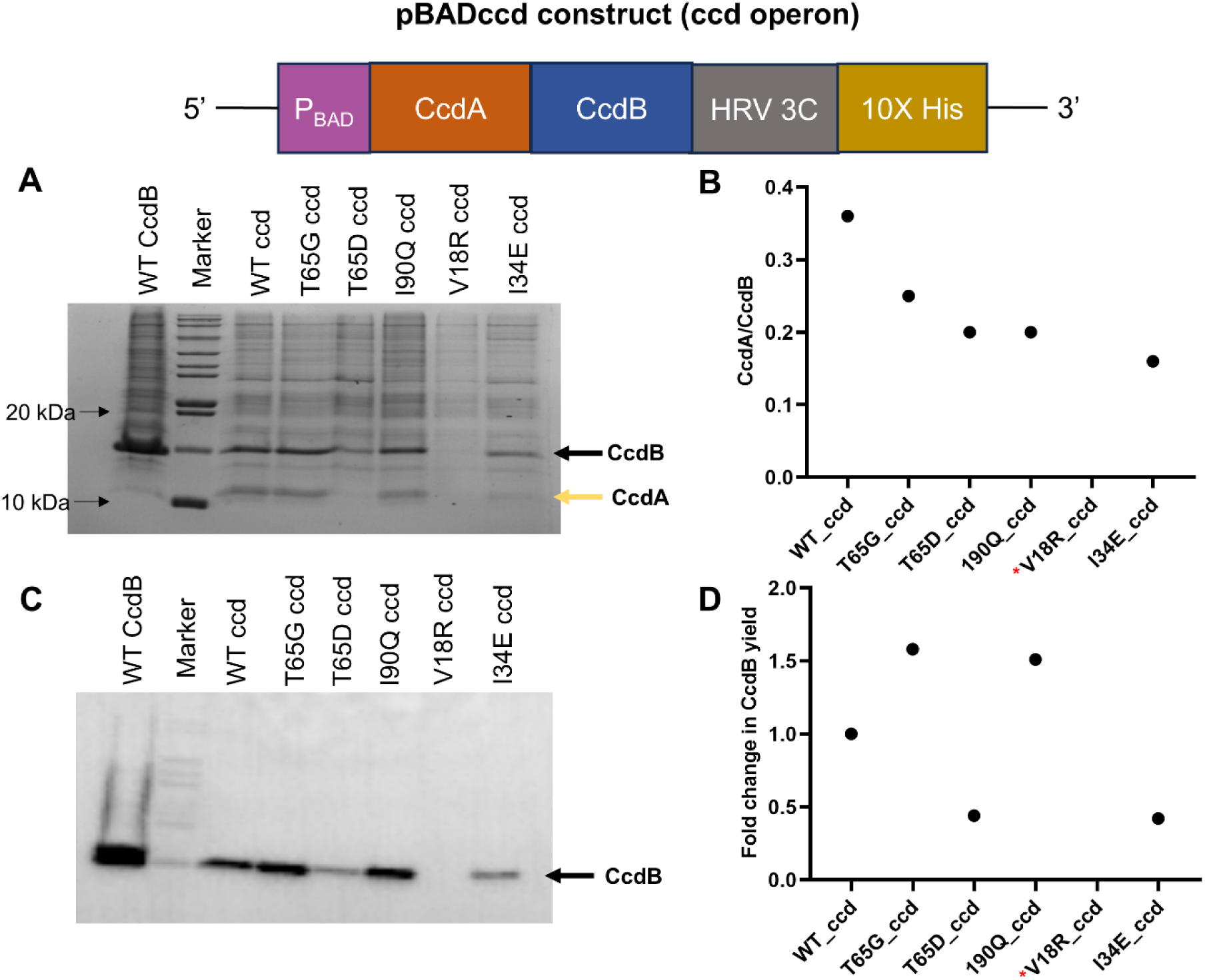
Comparison of protein yield of CcdB mutants when both CcdB and CcdA are expressed from the same polycistronic mRNA expressed in an operonic context. Top panel shows the construct design for the toxin-antitoxin arrangement in the pBAD vector where HRV 3C is the cleavage site of Human rhinovirus 3C protease. Positions of CcdB and CcdA bands are shown in black and yellow arrows, respectively. Final elutes were pooled and adjusted to the same volume. Equal volumes of the concentrated eluted protein were loaded and analyzed by (A) Coomassie stained SDS PAGE and (C) Western blot. (B) The ratio of copurified CcdA with CcdB was determined using SDS PAGE, and (D) fold change in CcdB protein yield for mutants relative to WT was determined using Western blotting (* represents no detectable band in the operonic context). CcdA and CcdB identities were confirmed by mass spectrometry (Figure S9) Some extra bands are also visible, which could either be chaperones or proteins that eluted non-specifically during Ni-NTA affinity purification.

### Structural analysis of relevant complexes

Structures of the (CcdB)_2_:CcdA(37–72)_2_ complex, and the CcdA (1–40) N terminal dimerization domain are shown in Figure 4. Buried surface areas involved in the various interfaces are listed in Table 1. From the structure of the (CcdB)_2_:CcdA(37–72)_2_ complex, it is clear that individual CcdA monomers in each CcdA dimer make contact with two different (CcdB)_2_ homodimers (see schematic in Figure 1S C). A large interface is formed between the C-terminal CcdB binding domain of CcdA and CcdB. In this complex, the CcdB binding domain of CcdA forms a substantially larger interface with one of the two monomers of the (CcdB)_2_ homodimer. This CcdB binding domain on CcdA is natively unfolded in the absence of CcdB and CcdA dimerization is not required for binding to CcdB (39, 52). In addition, the C-terminus of CcdB makes a substantial contribution to the (CcdB)_2_ homodimer interface, suggesting that dimerization of CcdB will likely occur after the chain is completely synthesised. We speculate that during protein synthesis, the natively unfolded C-terminal region of CcdA interacts with and stabilizes the CcdB nascent chain, prior to its homodimerization. If binding of CcdA to CcdB occurs early in the assembly process, this would also prevent interaction of CcdB with DNA Gyrase.

**Figure 4:**
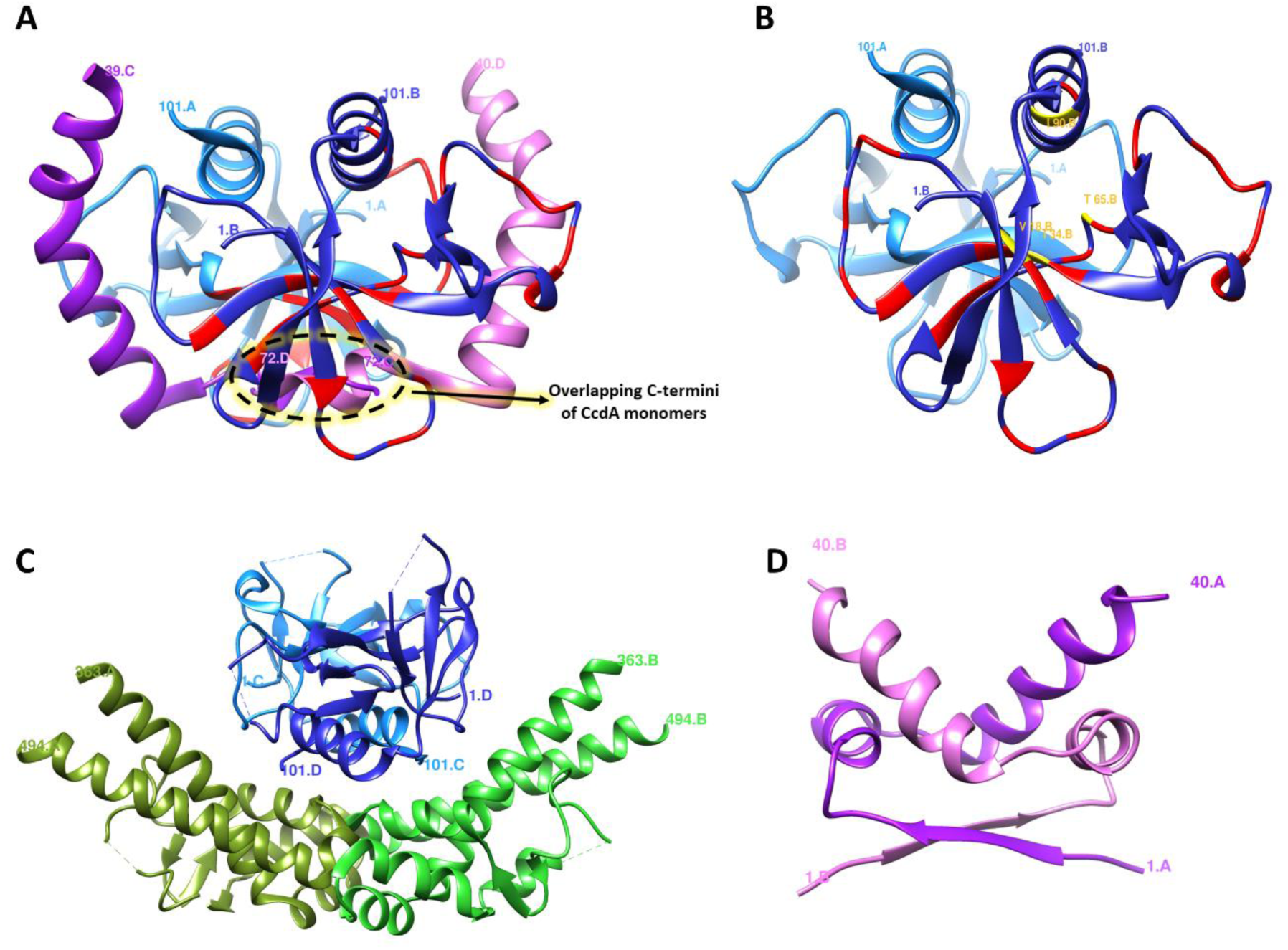
Structures of CcdA, CcdB and the CcdB:GyrA and CcdAB complexes. (A) CcdB:CcdA (37–72) complex: CcdB dimer is shown as a ribbon in different shades of blue, and the two C-terminal CcdA (37–72) peptides are shown as ribbons in purple and pink color. CcdB (chains A, B) residues interacting with CcdA (chain D) are shown in red (PDB-3G7Z). B) CcdB dimer: Two chains shown in different shades of blue (PDB-3G7Z). CcdB (chains A, B) residues interacting with CcdA (chain D) are shown in red. CcdB residue positions where mutants showed enhanced yield (see Figure 3) in the presence of operonic CcdA are shown in yellow. C) CcdB:Gyrase complex: CcdB dimer is shown as a ribbon in different shades of blue and GyraseA14 dimer is shown in different shades of green (PDB-1X75). D) CcdA (1–40) dimer: The N-terminal CcdA dimer is shown in purple and pink (PDB-2ADL).

**Table 1:**
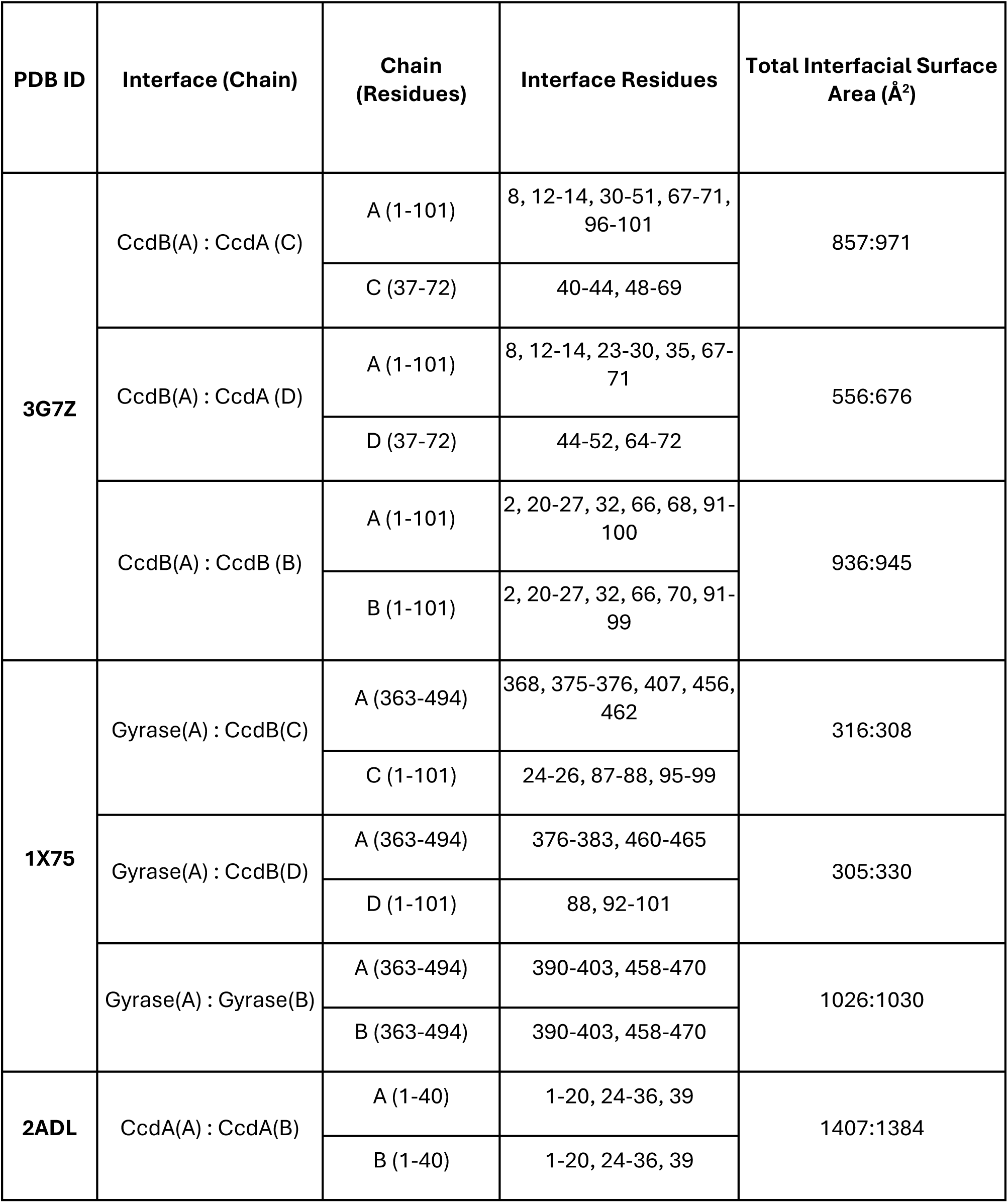
Details of Interfacial residues and total interface area of chains of CcdA:CcdB heterodimer, CcdB homodimer and N-terminus CcdA. (1–40) homodimer.

| PDB ID | Interface (Chain) | Chain (Residues) | Interface Residues | Total Interfacial Surface Area (Å<sup>2</sup>) |
| --- | --- | --- | --- | --- |
| **3G7Z** | CcdB(A) : CcdA (C) | A (1-101) | 8, 12-14, 30-51, 67-71, 96-101 | 857:971 |
|  |  | C (37-72) | 40-44, 48-69 |  |
|  | CcdB(A) : CcdA (D) | A (1-101) | 8, 12-14, 23-30, 35, 67-71 | 556:676 |
|  |  | D (37-72) | 44-52, 64-72 |  |
|  | CcdB(A) : CcdB (B) | A (1-101) | 2, 20-27, 32, 66, 68, 91-100 | 936:945 |
|  |  | B (1-101) | 2, 20-27, 32, 66, 70, 91-99 |  |
| **1X75** | Gyrase(A) : CcdB(C) | A (363-494) | 368, 375-376, 407, 456, 462 | 316:308 |
|  |  | C (1-101) | 24-26, 87-88, 95-99 |  |
|  | Gyrase(A) : CcdB(D) | A (363-494) | 376-383, 460-465 | 305:330 |
|  |  | D (1-101) | 88, 92-101 |  |
|  | Gyrase(A) : Gyrase(B) | A (363-494) | 390-403, 458-470 | 1026:1030 |
|  |  | B (363-494) | 390-403, 458-470 |  |
| **2ADL** | CcdA(A) : CcdA(B) | A (1-40) | 1-20, 24-36, 39 | 1407:1384 |
|  |  | B (1-40) | 1-20, 24-36, 39 |  |

## DISCUSSION

Several studies suggest that the folding trajectory of proteins while being translated by ribosomes can be altered via mechanisms involving cotranslational folding (24, 53, 54). The vectorial nature of translation is often defined by the early onset of folding events. Recent studies suggest that many proteins fold in a sequential, domain-wise fashion that helps to avoid kinetic traps of misfolding during an ongoing translation event on the ribosome (18, 27, 55). There are studies that suggest that due to sequential folding, synonymous mutations can alter the final folded form or yield of the protein (27, 28, 31, 32, 56–58). Apart from studies involving only synonymous substitutions, the importance of cotranslational assembly has also been illustrated in the efficiency of formation of stable protein complexes when both the interacting partners are synthesised from the same polycistronic mRNA (6). However, how cotranslational subunit assembly can help alleviate the folding defect of a misfolding or aggregation prone protein has not been well studied. Having interacting gene products expressed from the same operon is expected to promote assembly and it was shown that operon gene order correlates with predictions of the order in which gene products would assemble (4). A subsequent study (59) suggested that the largest interface in a complex corresponds to the earliest forming subcomplex and that N-terminal interfaces are on average larger than C-terminal interfaces.

In one of the few detailed experimental studies of cotranslational assembly in an operonic context, the Bukau group demonstrated that in the bacterial *Vibrio harveyi* luxAB operon that encodes the alpha and beta subunits of luciferase, assembly involves the cotranslational interaction of nascent LuxB with full-length and presumably folded LuxA. In the native operon, the two genes are separated by ∼26 bp (60). This study showed that gene organization within operons significantly affects the efficiency and mechanism of protein complex assembly, with subunits assembling more efficiently when expressed from the same operon. The methodology involved expression from a non-native Ptac promoter and included N-terminal fluorescent tags (YFP and CFP) fused to LuxA and LuxB, facilitating fluorescence resonance energy transfer (FRET) analysis. While this approach provided valuable insights into cotranslational assembly and operon dynamics, it also introduced large tags at the N-terminus of the genes, potentially affecting the translation efficiency of the component genes in the operon. In addition, in that study cotranslational assembly resulted in a relatively small enhancement of about two fold relative to expression from two different chromosomal locations. An earlier study (61) showed that untagged luciferase assembles with similar efficiencies when the two subunits are expressed from independent transcriptional units and when expressed in their native operonic context.

Here, we employed a site-saturation mutagenesis library of a globular cytotoxin, CcdB, both in the absence and presence of its interacting partner, CcdA. In contrast to the luxAB operon where the LuxAB heterodimeric complex consists of two well-folded globular proteins tagged at the N-terminus, the present ccdAB operonic system involves the binding of a natively unfolded segment of the antitoxin (CcdA) with the globular toxin (CcdB). In the native operon the Shine-Dalgarno sequence for the downstream ccdB gene is located just upstream of the stop codon of the ccdA gene. Our previous studies (31, 32) have shown that even single synonymous mutations, including at the junction between ccdA and ccdB can exhibit profound phenotypic effects. We have therefore entirely avoided the use of tags both at the N and C-termini of CcdA and the N-terminus of CcdB. The majority of the mutational phenotypes are observed in the native operonic context without any extraneous tags ensuring that the folding rescue and assembly processes are near physiological. Both CcdA and CcdB form stable homodimers in isolation, but form either heterohexameric (CcdB)_2_-(CcdA)_2_-(CcdB)_2_ or extended higher order (CcdB:CcdA)_n_ complexes, depending on the CcdA:CcdB ratio (39). Homodimeric CcdB is a highly toxic protein that binds and poisons its cellular target DNA Gyrase (62). Hence CcdA has to bind CcdB before it complexes with DNA Gyrase. CcdAB is a representative member of the ubiquitous family of bacterial Type II Toxin-Antitoxin systems. (39)(62)All of these features make this a very interesting system to characterize in terms of its assembly *in vivo.* We observed mutations present in a non-operonic context that resulted in an inactive phenotype due to protein misfolding, but displayed an active phenotype in an operonic context. *In vivo* studies reveal that several destabilised mutants of the CcdB toxin are rescued when expressed in an operonic context. These results suggest that CcdA traps mutated CcdB in a native-like form before it can aggregate, and then prevents it from unfolding due to a slow off-rate, implying that CcdA rescues folding defects in CcdB by enhancing kinetic stability *in vivo*. CcdB expression was not observed for mutants with low RF^RelE^ scores, suggesting that these CcdB proteins misfolded to such an extent that CcdA was not able to assist folding, and no complex was formed. Destabilised, unassembled subunits are cleared from the cytosol, this is termed as orphan protein degradation (13).

Comparison of the amount of induced CcdB WT and mutant protein found in the soluble fraction when the toxin and the antitoxin are expressed from separate mRNAs relative to when they are expressed from the same polycistronic mRNA suggests that CcdA can suppress folding defects of CcdB, likely via cotranslational assembly mechanisms. (Figure 2 B). We also observed that soluble protein expression in an operonic context is largely consistent with values of the RF^RelE^ scores obtained from the deep sequencing data.

Although the exact mechanism by which CcdA relieves the folding defect of CcdB still needs to be investigated, we speculate that the intrinsically disordered segment of CcdA acts as a chaperone, binding to nascent CcdB. Some intrinsically disordered proteins are known to act as highly specific steric chaperones (63). *In vitro* studies on another type II TA system Phd/Doc showed that the intrinsically disordered domain of Phd (antitoxin) binds to Doc (toxin) in its monomeric state, prevents domain swapping of Doc, stabilises it and prevents it from aggregation or misfolding (64). In the present study, it appears that cotranslational assembly is seeded during translation, the CcdB interacting, natively unfolded C-terminal domain of CcdA is recruited to the nascent CcdB chain preventing erroneous associations with other proteins including other toxins or antitoxins in the crowded cellular environment, thereby ensuring correct localisation and activity (1). Interestingly, although several non-cognate TA complexes can be formed *in vitro*, there appears to be a higher degree of specificity *in vivo*, since the toxicity seen on expression of a specific toxin often fails to be rescued by heterologous expression of a non-cognate antitoxin that is able to form a stable complex with the toxin *in vitro* (65). The Shine-Dalgarno sequence for the *ccdB* component is located towards the C-terminal end of the *ccdA* gene, before the stop codon. Future studies will explore the significance of this.

Several other studies in eukaryotes have also shown that cotranslational assembly is indeed advantageous in preventing aggregation or degradation (55, 66–68). The above-mentioned studies involve an oligomeric complex wherein the subunit genes are positioned at distinct locations on the chromosome. This might decrease the efficiency of complex formation, though several studies have now shown that protein:protein interactions during translation can facilitate co-localization of heterologous mRNAs and promote co-translational assembly (69, 70). In the present study, both the CcdA and CcdB interacting partners are part of the same polycistronic mRNA. We show how an inherent folding defect in one of the partners, due to the presence of a non-synonymous mutation in the protein core, is rescued in the presence of its cognate partner. In Type II TA systems, the toxin component has a relatively conserved fold. Although the toxin binding regions on the antitoxin are helical, the binding interfaces are poorly conserved in different Type II TA complexes.

Taken together, these insights suggest cotranslational assembly is intimately linked with various cellular phenomena including regulation of protein abundance, kinetic stability and activity, and orphan subunit degradation. Relative rescue of folding-defective mutations in an operonic context compared to when the interacting partner is located on an independently transcribed mRNA, provides a convenient tool to assess whether co-translational assembly is occurring, and the present approach can readily be applied to other Type II TA systems as well as other operons. Gene organization of interacting partners in an operon allows for rescue of folding defects, thus providing a buffering mechanism to enhance genetic variation, and facilitating evolution of complexes in which toxin and antitoxin bind each other in a variety of different geometries and quaternary structures (71).

## EXPERIMENTAL PROCEDURES

### Plasmids and Host Strains

A site-saturation mutagenesis library containing ∼1600 single-site mutants of CcdB present in a non-operonic context in pBAD24 vector (44) and also in an operonic context in pUC57 vector (41) is used in this study. Top10Gyr and BL21(DE3)pLysE *E. coli* host strains are used for expression and purification of WT and CcdB mutants. A few *ccdB* mutants with and without *ccdA* are cloned in an arabinose inducible pBAD24 vector. WT *ccdB* and a few *ccdB* mutants are cloned with a cleavable C-terminal His-tag in the operonic context in pBAD24 vector. WT *ccdB* and a few *ccdB* mutants, along with WT *ccdA* are cloned under separate but identical T7 promoters in pET-Duet-1 vector. The tZ terminator (51) was inserted 32 bp downstream of the ccdB stop codon by Gibson assembly.

### Deep sequencing analyses

Deep sequencing of a site-saturation mutagenesis library of CcdB alone, i.e., in the absence of CcdA has been previously carried out. The site-saturation mutagenesis library generated in the absence of CcdA was plated on increasing arabinose (inducer) concentration and decreasing (repressing) glucose concentration. WT CcdB showed an active phenotype and kills cells at the lowest expression level, i.e., MID 2. The activity of several other mutants can be estimated from their growth as a function of the expression level in terms of their MID. Variant scores in the form of MS_seq_ values were assigned to each mutant which is defined as the MID at which the number of reads for a particular mutant decreases by 5-fold or more relative to its previous MID. These ranged from 2-9 with two showing a WT like phenotype and in increasing order of inactive phenotype relative to WT (43, 44). Deep sequencing of the same library in the operonic context has also been carried out previously. Read numbers for all mutants at all 101 positions (1–101) in CcdB were analysed.

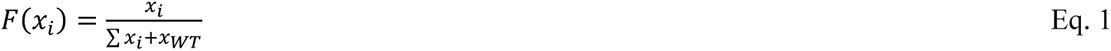

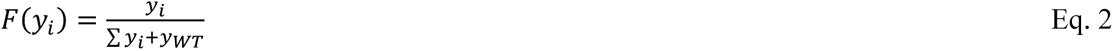

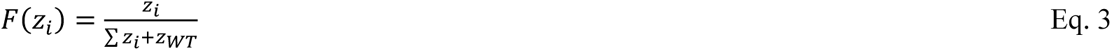

Here, a given mutant is represented by ‘i’ whereas WT is represented by ‘WT’. Number of reads in Top10Gyr resistant strain, Top10 sensitive strain and RelE reporter strain is represented by ‘x’, ‘y’ and ‘z’, respectively. F(x_i_), F(y_i_) and F(z_i_) are the fraction representation of a mutant in resistant, sensitive and RelE reporter strain, respectively.

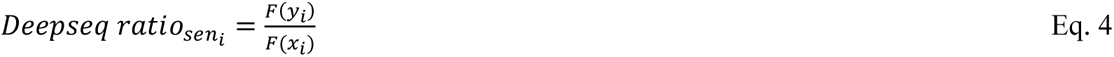

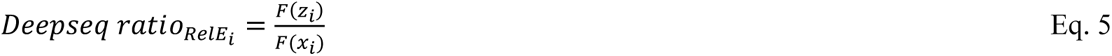

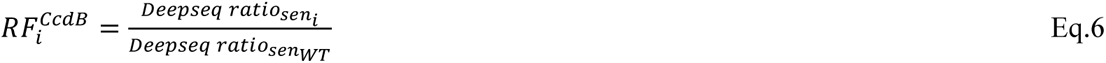

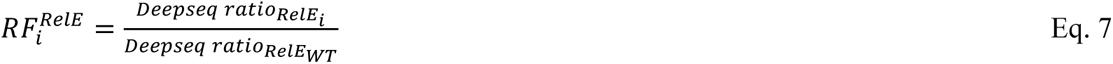

For simplicity, these mutational scores are represented as RF^CcdB^ and RF^RelE^ for each mutant throughout the text. RF^CcdB^ is based on the CcdB toxicity readout in the Top10 strain (CcdB sensitive strain) while RF^RelE^ is based on the RelE toxicity readout in Top10Gyr strain harbouring the RelE reporter gene (RelE reporter strain). The two variant scores are generally indicated in linear scale throughout the text. WT scores for RF^CcdB^ and RF^RelE^ are 1 (41).

z-scores of all the mutants and WT for each parameter were calculated.

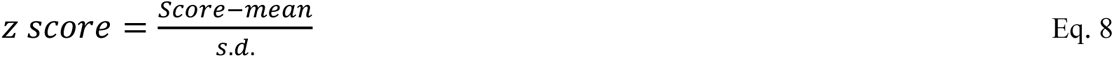

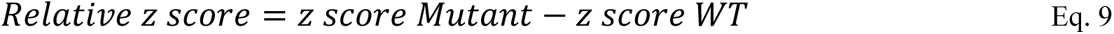

### Basis of classification of CcdB mutants into different structural categories

Depth and accessibility of each residue in CcdB was calculated based on the available structure from Protein Data Bank for dimeric CcdB structure (PDB ID: 3VUB) (72). The depth calculation is carried out using DEPTH server. In DEPTH server, depth is defined as the distance of any atom/residue to the closest bulk water. (73–75). A probe size of 1.4 Å was used to calculate accessibility. A residue was defined as buried or exposed if the side chain accessibility is ≤5% or >5%, respectively, based on the accessibility calculated using NACCESS program (41, 43, 76). Based on these calculations, CcdB mutants were segregated into different structural categories as described previously (41).

### Expression and *in-vivo* solubility of CcdB mutants

A subset of aggregation-prone CcdB mutants (V18R, D19M, I34E, I34R, L36D, L36R, T65D, T65G, L83Y, I90Q, I94L and F98R) were cloned alone as well as downstream of WT CcdA in an operonic context in pBAD24 vector by Genscript, USA. The corresponding proteins were heterologously expressed in pBAD24 vector in Top10Gyr *E.coli* strain that is resistant to the toxic action of CcdB. Expression levels were monitored for these CcdB mutants in *E. coli* Top10Gyr strain in the presence of 0.2% arabinose. Mutants were grown in LB medium, induced with 0.2% arabinose at an OD_600_ of 0.6, and grown for 4.5 hours at 37 °C/180 rpm. Equal numbers of cells (2.4X10^9^ cells) for all the CcdB variant proteins were taken and centrifuged at 4000 rpm for 10 min at room temperature. The pellet was resuspended in resuspension buffer (pH 7.4) and sonicated, followed by centrifugation at 12700 rpm for 10 min at 4 °C. The supernatant and pellet were separated, individually made up to 60 μL in 1× loading buffer and subjected to 15% Tricine sodium dodecyl sulfate−polyacrylamide gel electrophoresis (Tricine SDS−PAGE). Following staining with Coomassie Blue R-250, the solubility of all mutants was estimated by quantitating the relative amounts in supernatant and pellet fractions using the geldoc software Quantity One (Bio-Rad). A few CcdB mutants (V18R, I34E, T65G, T65D and I90Q) along with the WT were also cloned in pET-Duet-1 vector. These mutants were transformed in BL21(DE3)pLysE *E.coli* strain to co-express toxin and antitoxin from separate promoters. Here CcdB and CcdA were induced by 1mM IPTG. The same protocol as described above was used.

### Purification of CcdAB complex and CcdB protein

WT and five destabilised CcdB mutant complexes (V18Rccd, I34Eccd, T65Gccd, T65Dccd and I90Qccd) cloned in pBAD24 vector were purified using Ni-NTA affinity chromatography. These variants contained a C-terminal His tag and were expressed in *E. coli* Top10Gyr strain. A single colony was inoculated in 5 mL LB medium containing 100 µg/μL ampicillin and grown at 37 °C overnight. 1% of the primary inoculum was inoculated in 500 mL of secondary culture in LB media containing 100 µg/μL ampicillin. Cells were grown to an OD_600_ of 0.6 at 37 °C in LB medium, induced with 0.2% (w/v) arabinose, and grown at 37 °C for 4.5 hours. Cells were harvested by centrifuging at 4000 rpm at 4 °C for 15 min, resuspended in 20 mL resuspension buffer (10 mM HEPES + 100 mM NaCl + 100 mM Arginine + 10% Glycerol + Roche Protease Inhibitor cocktail), sonicated on ice, and centrifuged at 14000 rpm, 4 °C for 30 min. The supernatant was loaded onto a Ni-NTA affinity column packed with Ni-Sepharose Fast Flow beads and incubated at 4 °C for 2-3 hours. The column was further washed with 50-60 ml of wash buffer (10 mM HEPES + 100 mM NaCl + 100 mM Arginine + 50 mM Imidazole). Elutes were carried out with increasing concentrations of Imidazole in elution buffer (10 mM HEPES + 100 mM NaCl + 100 mM Arginine + 100-1000 mM Imidazole). The eluted fractions were concentrated and stored at −80 °C. The eluted fractions were subjected to 15% Tricine SDS−PAGE to estimate protein yield relative to WT.

WT CcdB with a cleavable C-terminal His-tag was cloned in pBAD24 vector in Top10Gyr *E.coli* strain. The same protocol was followed as described above.

### Western blotting

After concentrating and adjusting the purified proteins containing a C-terminal His-tag to equal volume, they were run on 15% Tricine SDS-PAGE. Following SDS-PAGE, proteins were electrophoretically transferred to a polyvinylidene difluoride membrane (Millipore). The membrane was blocked with 5% skim milk for 1 hour. The membrane was washed with PBST (PBS with 0.05 % Tween) and incubated with Anti-His IgG conjugated with Horseradish peroxidase (Sigma) in PBS at 1:10,000 dilution, for 1 hour. The membrane was washed again with PBST, and bound antibody was detected with chemiluminescence using HRP substrate and luminol in a 1:1 ratio (Bio-Rad). Protein band intensities were measured via densitometric analysis of Western blot data using the Image Lab software from Bio-Rad.

## DATA AVAILABILITY

The variant scores (MS_seq_, RF^CcdB^ and RF^RelE^) obtained from deep sequencing have been taken from previous reports (41, 44). The raw deep sequencing data used in the present study has been deposited in NCBI’s Sequence Read Archive (accession no. SRR17982061). Remaining data are available in the manuscript.

## ACKNOWLEDGEMENTS

PB acknowledges University Grants Commission, Government of India, for her fellowship. PK acknowledges Ministry of Human Resource and Development, Government of India, for her fellowship. RV is a J. C. Bose Fellow of DST.

## AUTHOR CONTRIBUTION

R.V. conceptualization; P.B., P.K. and R.V. methodology; P.B. and P.K. validation; P.B. formal analysis; P.B. and P.K. investigation; P.B. and P.K. data curation; P.B. and P.K. software; P.B. and P.K. visualization; R.V. resources; P.B. and R.V. writing—original draft; P.B., P.K. and R.V. writing - review & editing; R.V. supervision; R.V. project administration; R.V. funding acquisition.

## SUPPORTING INFORMATION

This article contains supporting information.

**Table S1:**
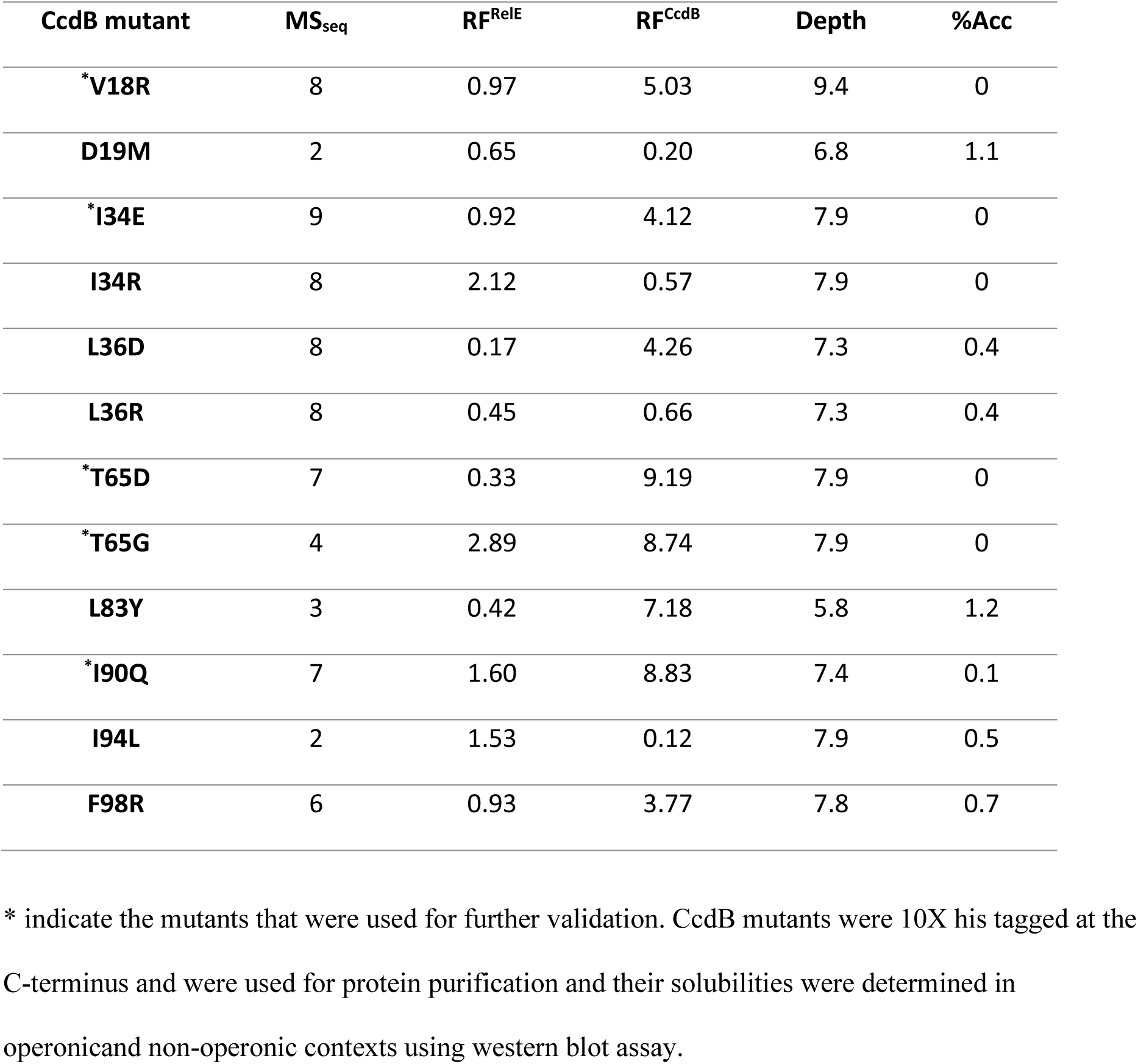
Details of the mutants used for validating deep sequencing results via *in vivo* solubility assays when expressed either alone or in an operonic context.

**Figure S1:**
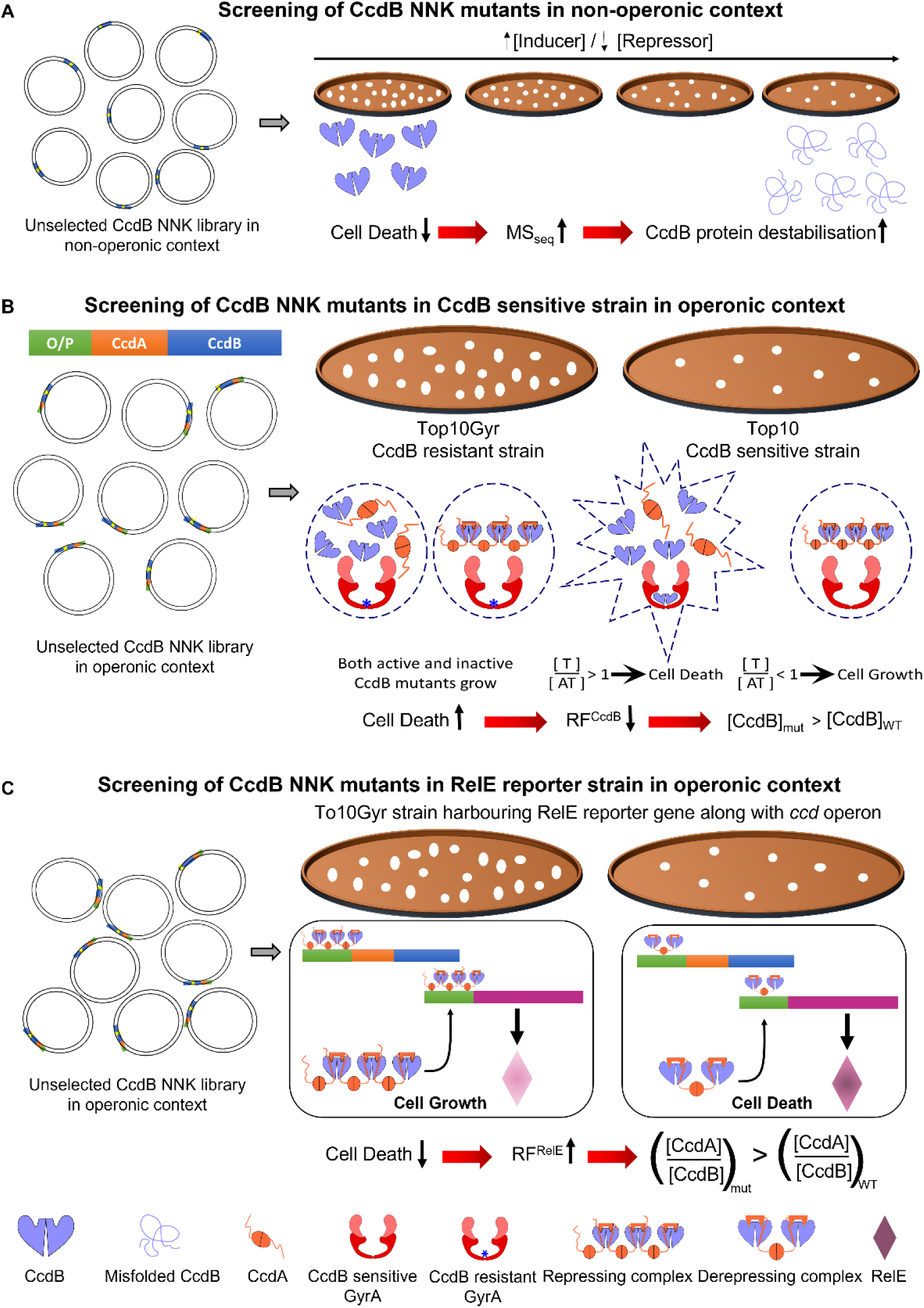
Pictorial depiction of the strategies for the three phenotypic readouts for which the deep sequencing data is inferred. (A) Schematic for screening of CcdB NNK mutants in a non-operonic context. Here, CcdB is expressed under the control of arabinose inducible promoter. The CcdB NNK library is plated on increasing arabinose (inducer) and decreasing glucose (repressor) concentrations. If the mutation is in the protein core then higher MS_seq_ values are correlated with enhanced destabilisation of the protein (1). (B) Schematic of screening of CcdB NNK mutants in an operonic context in CcdB sensitive and resistant strains. The CcdB resistant strain Top10Gyr, contains an R462C mutation in GyrA that prevents CcdB from binding to DNA Gyrase (shown by a blue asterisk mark in the GyrA subunit). The CcdB NNK library prepared in the CcdB resistant strain was transformed into the CcdB sensitive strain to screen for active versus inactive mutants in terms of Gyrase poisoning/inhibition activity. RF^CcdB^ scores are greater than 1 for mutants with decreased activity against Gyrase (2). (C) Schematic for screening of CcdB NNK mutants in operonic context in RelE reporter strain. The CcdB NNK library (2) was transformed in Top10Gyr strain harbouring the RelE reporter gene in pBT vector under the *ccd* promoter. This screen helps to understand the cause for the inactive phenotype observed for several buried-site CcdB mutants. For instance, an inactive and repressing phenotype implies that the CcdB structure is intact yet it fails to bind GyrA (Gyrase A subunit) likely because of mutations in the GyrA binding site. However, the mutant still forms a tight complex with CcdA which in turn represses RelE expression, thereby diminishing RelE toxicity. On the contrary, an inactive and derepressing phenotype implies that the CcdB structure is disrupted because of which both Gyrase as well as CcdA binding are abrogated.

**Figure S2:**
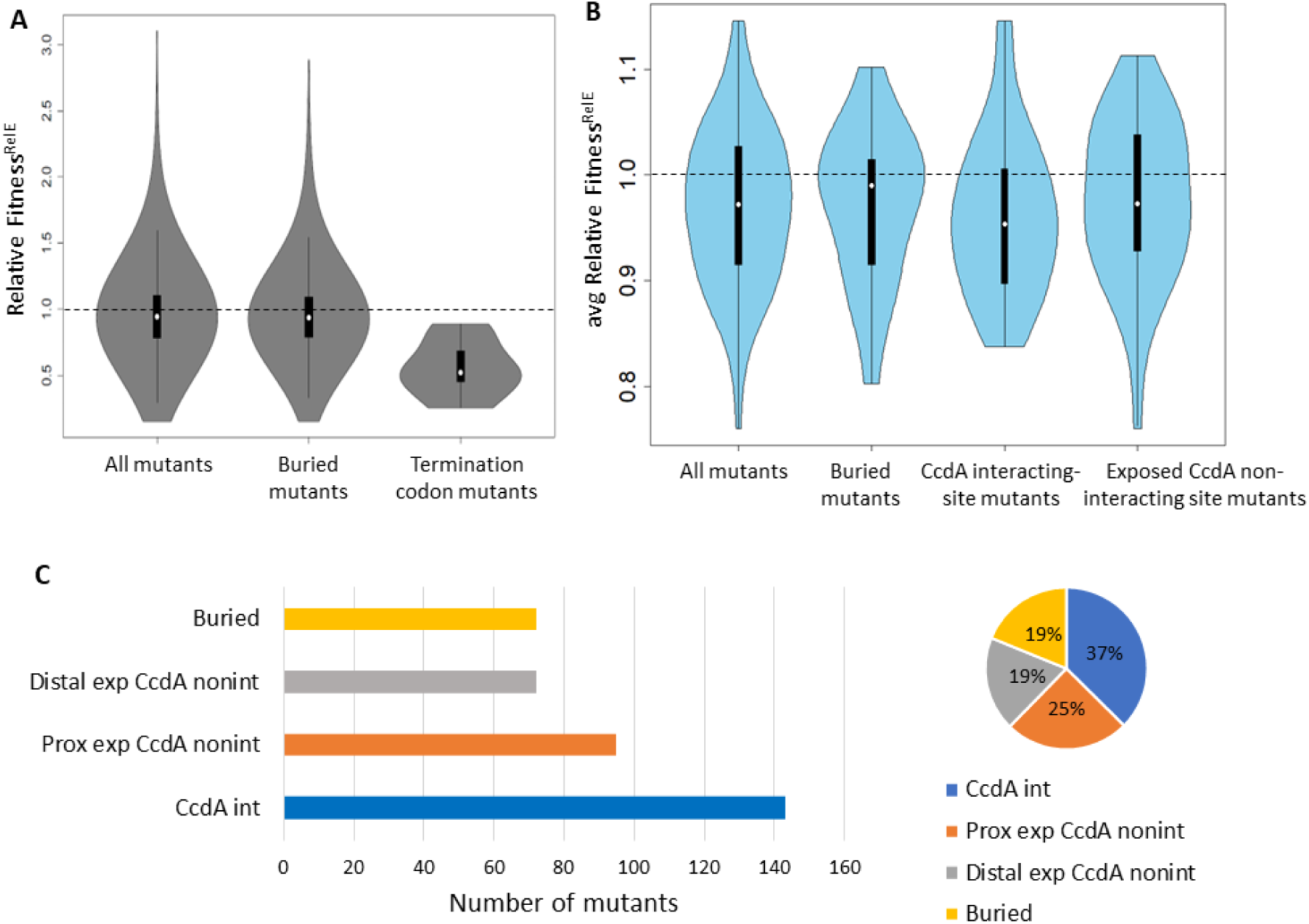
The majority of CcdB mutants in the protein core show an active phenotype when expressed in an operonic context. (A) Frequency distribution of RF^RelE^ for the entire dataset, for buried site mutants and TAA termination codon mutants. The majority of the buried-site mutants show a phenotype similar to the WT while the highest frequency stop codon in the *E.coli* genome, TAA gave the expected derepressing phenotype. (B) Distribution of RF^RelE^ scores averaged over each position for four structural categories, namely all mutants, buried site mutants, CcdA interacting site mutants, and exposed CcdA noninteracting site mutants. In (A-B), the width of the violin plot is proportional to the relative number of mutants. The white dot in the middle represents the median of the distribution. The box plot in black colour represents the inter-quartile range. The black dashed line corresponds to the normalised score obtained by the WT (RF^RelE^ = 1). (C) CcdA binding defective mutants with RF^RelE^<0.7, classified into four different structural classes, i.e., at the CcdA binding site (CcdA int), exposed and proximal (within 8Å) to the CcdA binding site (Prox exp CcdA nonint), exposed but distal (>8Å) to the CcdA binding site (Distal exp CcdA nonint) and in the protein interior (Buried). The length of the bar is proportional to the total number of mutants for each class (left panel). These absolute numbers are converted into percent fractions shown in a pie chart (right panel)

**Figure S3:**
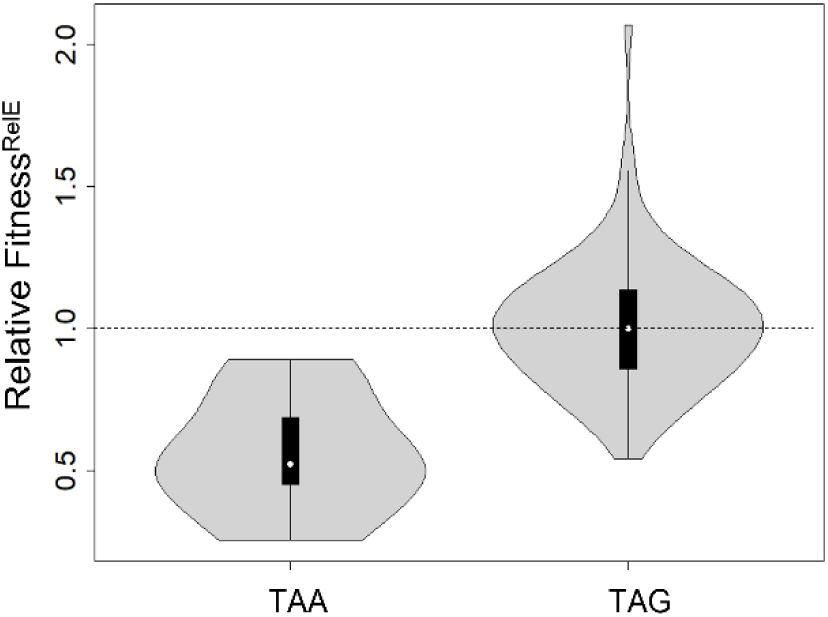
Frequency distributions of RF^RelE^ for two different types of nonsense mutants (TAA and TAG). The high frequency TAA stop codon mutant (also shown in Figure S2A) gave a derepressing phenotype whereas the TAG stop codon mutant spanned the entire range of scores for reasons not clearly understood.

**Figure S4:**
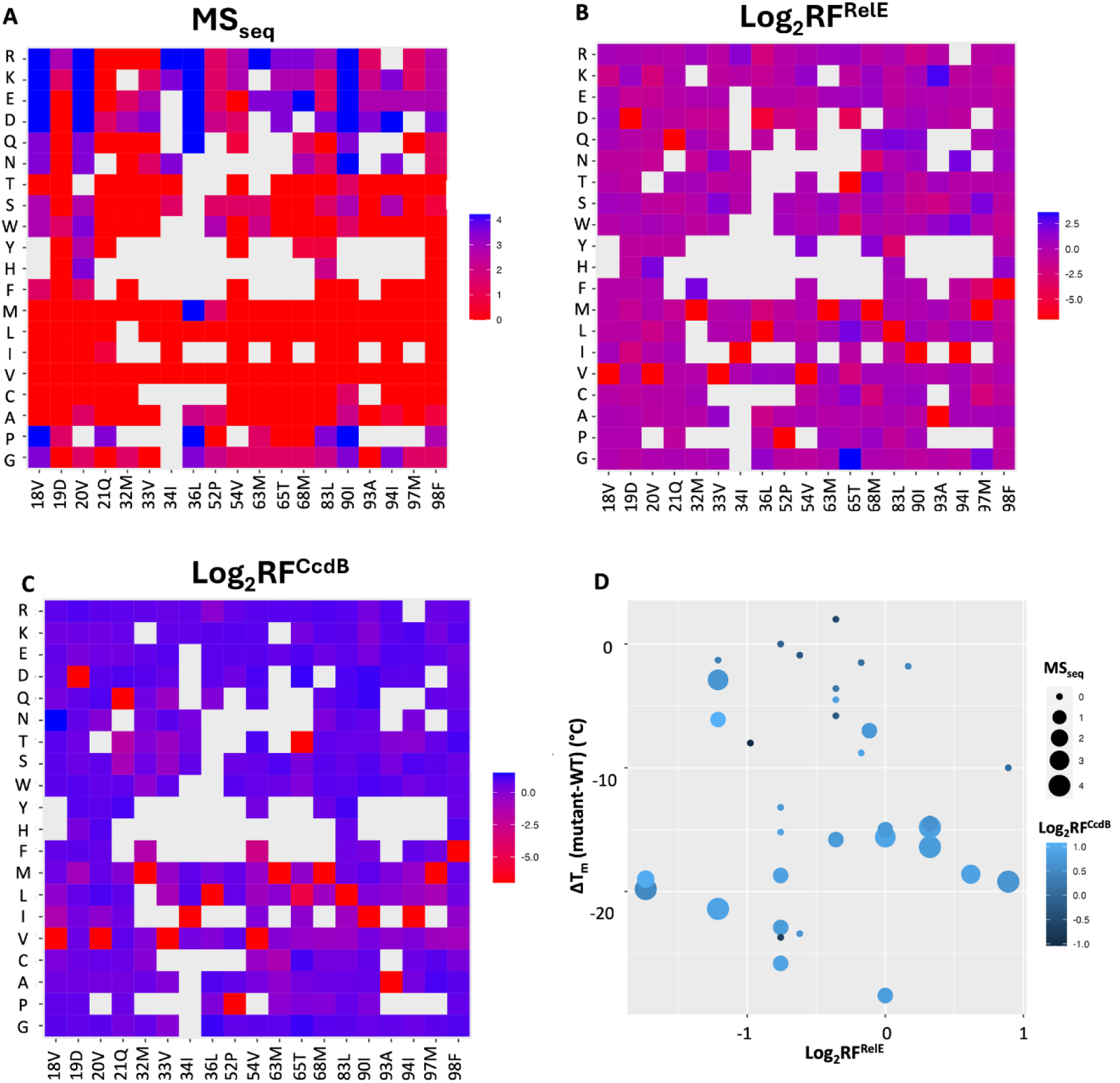
Mutational effects on CcdB activity inferred from phenotypic screening and deep sequencing, of CcdB buried-site mutants in the absence and presence of CcdA. MS_seq_, RF^RelE^ and RF^CcdB^ scores obtained from deep sequencing are averaged over synonymous mutants. For comparison, mutational scores were converted to z score and further represented relative to the WT z-score value. (A-C) On the left vertical axis, residues are grouped into (G, P), aliphatic (A–M), aromatic (F–W), polar (S–Q) and charged (D–R) amino acids. Residue numbers with their corresponding WT amino acid and substitutions are indicated on the horizontal and left vertical axes, respectively. Mutations where there is no data are shown in light grey. The colour scheme for the heat map is shown on the right of each plot, and the colour bar represents the relative z-scores. Relative z score for (A) MS_seq_ values for buried site mutants. Gradation of red to blue colour represents increasing relative z scores and decreasing CcdB activity. WT residue at each position is indicated in red, (B) RF^RelE^ scores for buried site mutants. Red to blue gradation represents increasing relative z scores, indicating increasing amounts of CcdAB complex. The WT residue at each position is indicated in red, (C) RF^CcdB^ for buried site mutants. A relative z score could either be because CcdB is neutralised by CcdA or it remains misfolded even in the operonic context. Gradation of red to blue corresponds to increasing relative z score. WT residue at each position is indicated in red colour. (D) Thermal stabilities of characterised buried-site mutants (3) were compared with their corresponding relative RF^RelE^, MS_seq_ and RF^CcdB^ scores. Coordinates (x,y) of each point reflect the corresponding relative RF^RelE^ value and the difference in thermal stability of the mutant with respect to WT (ΔT_m_). The size of the bubble increases with increasing MS_seq_ score. The blue colour becomes lighter with increasing RF^CcdB^ score. Smaller dots (lower MS_seq_ score) are typically darker (lower RF^CcdB^ score). Several highly destabilized mutants (ΔT_m_<-10°C) have RF^RelE^>1, indicating that they are well folded when expressed in the operonic context.

**Figure S5:**
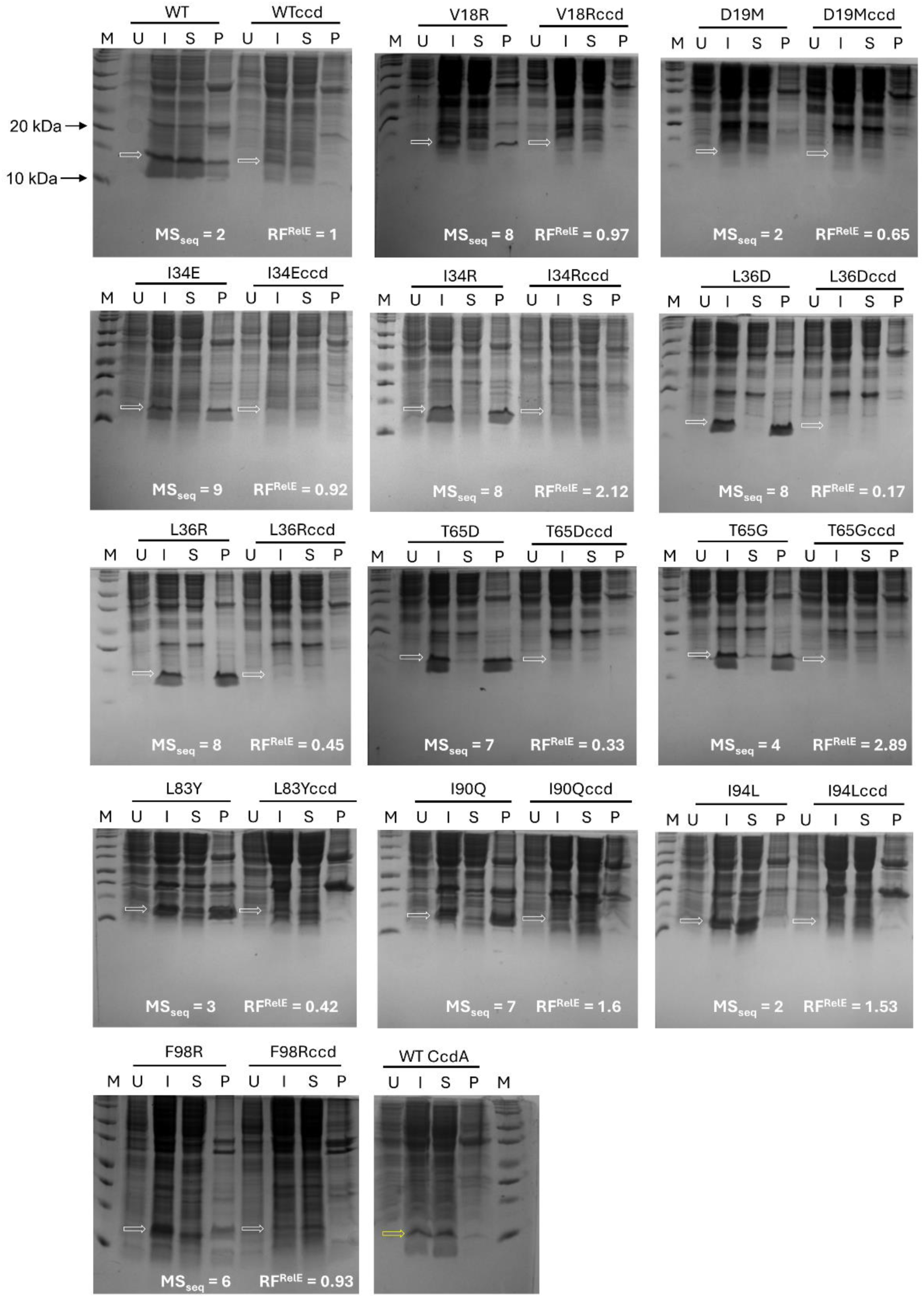
Expression of buried-site CcdB mutants in the absence and presence of CcdA. *In vivo* solubility of destabilised CcdB mutants when expressed alone relative to when they are present in an operonic context from an arabinose inducible promoter. The corresponding mutational score obtained from deep sequencing, i.e., MS_seq_ (-CcdA) and RF^RelE^ (+CcdA), for each mutant is mentioned on the gel. M represents the protein ladder. U, I, S and P represent uninduced, induced, supernatant and pellet cell lysate fractions, respectively. The fraction of protein in supernatant or pellet was determined by densitometric analysis following 15% Tricine SDS-PAGE and Coomassie staining. The white arrow indicates the likely position of the protein of interest (CcdB). The yellow arrow indicates the position of WT CcdA protein when expressed separately from an IPTG inducible promoter (bottom last panel). This assay was carried out in biological duplicates.

**Figure S6:**
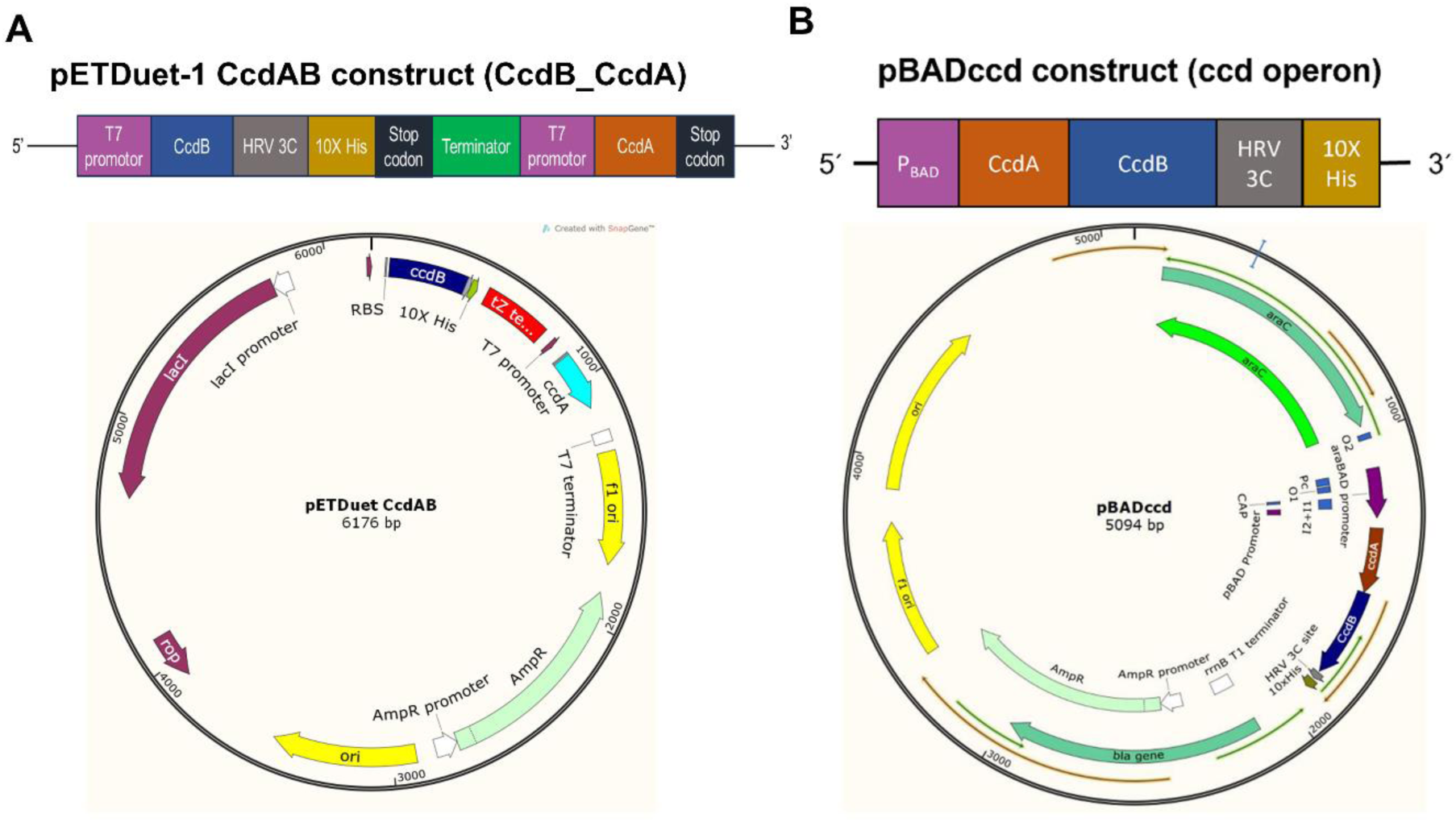
Two different vector systems were used to perform *in vivo* solubility assays and for purifying proteins. (A) pETDuet construct. Top panel shows *ccdB* and *ccdA* gene organization in the pETDuet vector. Here *ccdB* and *ccdA* are expressed separately under two different IPTG inducible T7 promoters in the same vector. The *ccdB* gene sequence is identical to that used in the pBADccd construct. Bottom panel shows the vector map of the construct. (B) pBADccd construct. Top panel shows gene organization of the *ccdAB* operon with a cleavable C-terminal His tag containing an HRV3C protease site in pBAD24 vector. ccdA and ccdB induction is under control of the arabinose inducible P_BAD_ promoter. Bottom panel shos vector map of the construct.

**Figure S7:**
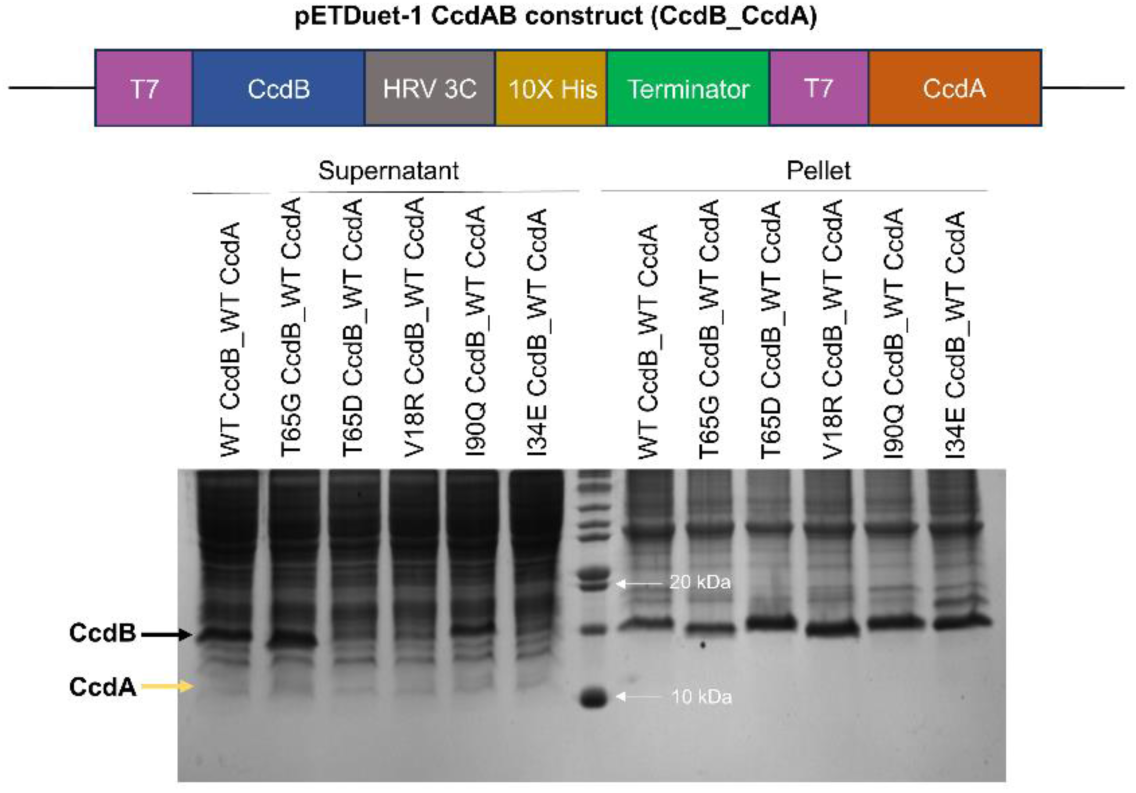
Coomasie stained SDS-PAGE of His-tagged CcdB mutants in the presence of CcdA, expressed from different mRNAs. *In vivo* solubility of Wild-type and destabilized CcdB mutants when expressed in a non-operonic context. Both CcdB and CcdA are expressed from two different T7 promoters. Black arrow indicates the CcdB protein and yellow arrow indicate CcdA protein. In order to visualize the CcdA bands, larger amounts of sample were loaded in this gel relative to those in Figure 2B.

**Figure S8:**
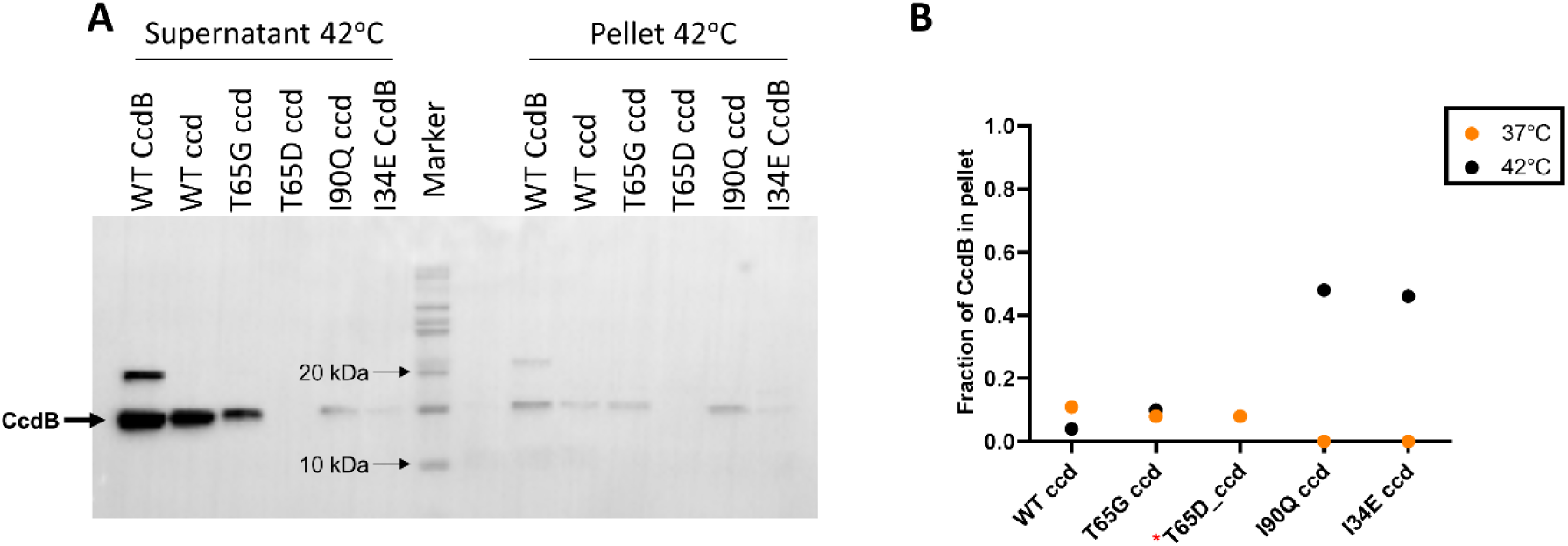
Western blot of His-tagged CcdB mutants in the presence of CcdA. *In vivo* solubility of Wild-type and destabilized CcdB mutants when expressed in an operonic context from an arabinose inducible promoter (A) at 42^ᵒ^C. (B) Comparison of fraction of CcdB in pellet relative to supernatant at 37^ᵒ^C and 42^ᵒ^C. The arrow indicates the protein of interest (CcdB). Data for 37^ᵒ^C are shown in Figure 2A. (* represents no detectable band in the operonic context at 42°C)

**Figure S9:**
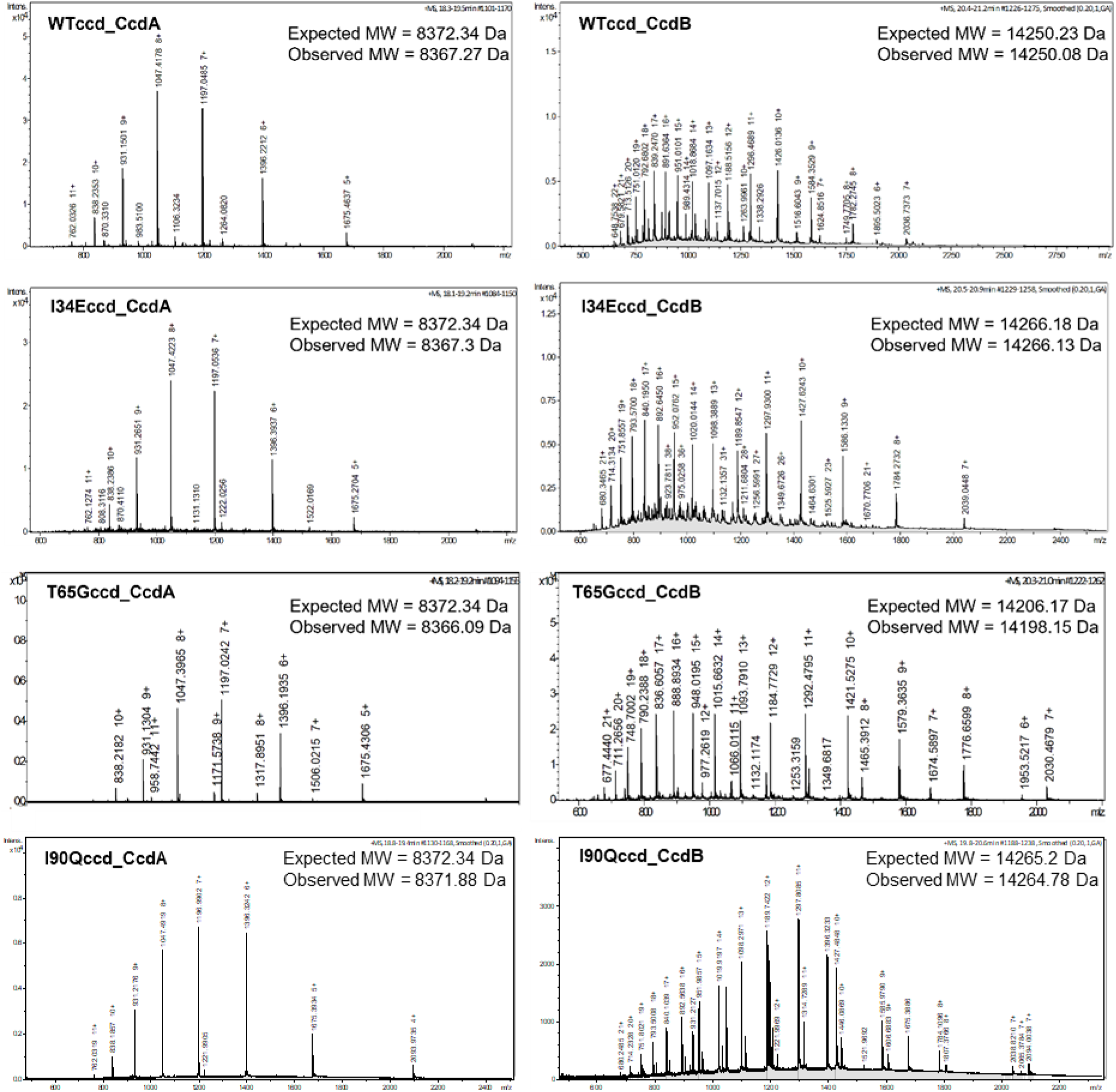
Mass spectrometry analysis of purified WT and mutant complexes when CcdA and CcdB are expressed in an operonic context. ESI Mass spectra of CcdA and CcdB protein components from mass spectrometry of CcdAB complex for WT and CcdB mutant (I34E, T65G and I90Q) proteins. Spectra for V18R could not be recorded because it could not be purified in sufficient amount due to its poor expression. In all cases the CcdB has a C-terminal His tag. Expected and observed <u>M</u>olecular <u>W</u>eights (MW) for the CcdB and CcdA components are mentioned above the spectra.

