## Supplementary material for "CcdA chaperones CcdB against irreversible misfolding and aggregation via a cotranslational folding mechanism": Table S1, Figure S1, Figure S2, Figure S3, Figure S4, Figure S5, Figure S6, Figure S7, Figure S8, Figure S9

**SUPPORTING INFORMATION**

**Table S1: Details of the mutants used for validating deep sequencing results via *in vivo* solubility assays when expressed either alone or in an operonic context.**

| CcdB mutant | MS_seq_ | RF^RelE^ | RF^CcdB^ | Depth | %Acc |
| --- | --- | --- | --- | --- | --- |
| ^*^V18R | 8 | 0.97 | 5.03 | 9.4 | 0 |
| D19M | 2 | 0.65 | 0.20 | 6.8 | 1.1 |
| ^*^I34E | 9 | 0.92 | 4.12 | 7.9 | 0 |
| I34R | 8 | 2.12 | 0.57 | 7.9 | 0 |
| L36D | 8 | 0.17 | 4.26 | 7.3 | 0.4 |
| L36R | 8 | 0.45 | 0.66 | 7.3 | 0.4 |
| ^*^T65D | 7 | 0.33 | 9.19 | 7.9 | 0 |
| ^*^T65G | 4 | 2.89 | 8.74 | 7.9 | 0 |
| L83Y | 3 | 0.42 | 7.18 | 5.8 | 1.2 |
| ^*^I90Q | 7 | 1.60 | 8.83 | 7.4 | 0.1 |
| I94L | 2 | 1.53 | 0.12 | 7.9 | 0.5 |
| F98R | 6 | 0.93 | 3.77 | 7.8 | 0.7 |

* indicate the mutants that were used for further validation. CcdB mutants were 10X his tagged at the C-terminus and were used for protein purification and their solubilities were determined in operonicand non-operonic contexts using western blot assay.

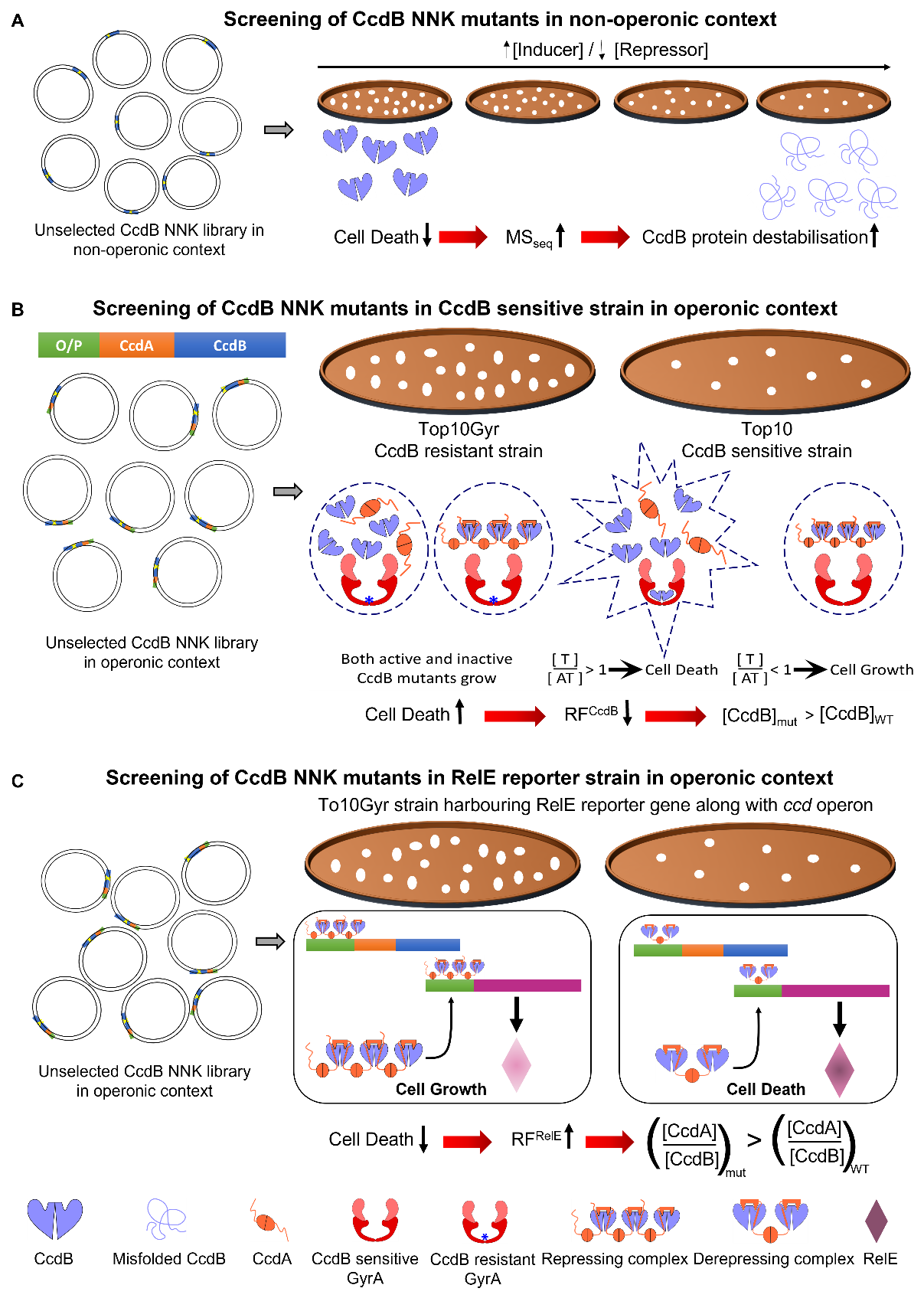

**Figure S1: Pictorial depiction of the strategies for the three phenotypic readouts for which the deep sequencing data is inferred.** (A) Schematic for screening of CcdB NNK mutants in a non-operonic context. Here, CcdB is expressed under the control of arabinose inducible promoter. The CcdB NNK library is plated on increasing arabinose (inducer) and decreasing glucose (repressor) concentrations. If the mutation is in the protein core then higher MS_seq_ values are correlated with enhanced destabilisation of the protein (1). (B) Schematic of screening of CcdB NNK mutants in an operonic context in CcdB sensitive and resistant strains. The CcdB resistant strain Top10Gyr, contains an R462C mutation in GyrA that prevents CcdB from binding to DNA Gyrase (shown by a blue asterisk mark in the GyrA subunit). The CcdB NNK library prepared in the CcdB resistant strain was transformed into the CcdB sensitive strain to screen for active versus inactive mutants in terms of Gyrase poisoning/inhibition activity. RF^CcdB^ scores are greater than 1 for mutants with decreased activity against Gyrase (2). (C) Schematic for screening of CcdB NNK mutants in operonic context in RelE reporter strain. The CcdB NNK library (2) was transformed in Top10Gyr strain harbouring the RelE reporter gene in pBT vector under the *ccd* promoter. This screen helps to understand the cause for the inactive phenotype observed for several buried-site CcdB mutants. For instance, an inactive and repressing phenotype implies that the CcdB structure is intact yet it fails to bind GyrA (Gyrase A subunit) likely because of mutations in the GyrA binding site. However, the mutant still forms a tight complex with CcdA which in turn represses RelE expression, thereby diminishing RelE toxicity. On the contrary, an inactive and derepressing phenotype implies that the CcdB structure is disrupted because of which both Gyrase as well as CcdA binding are abrogated.

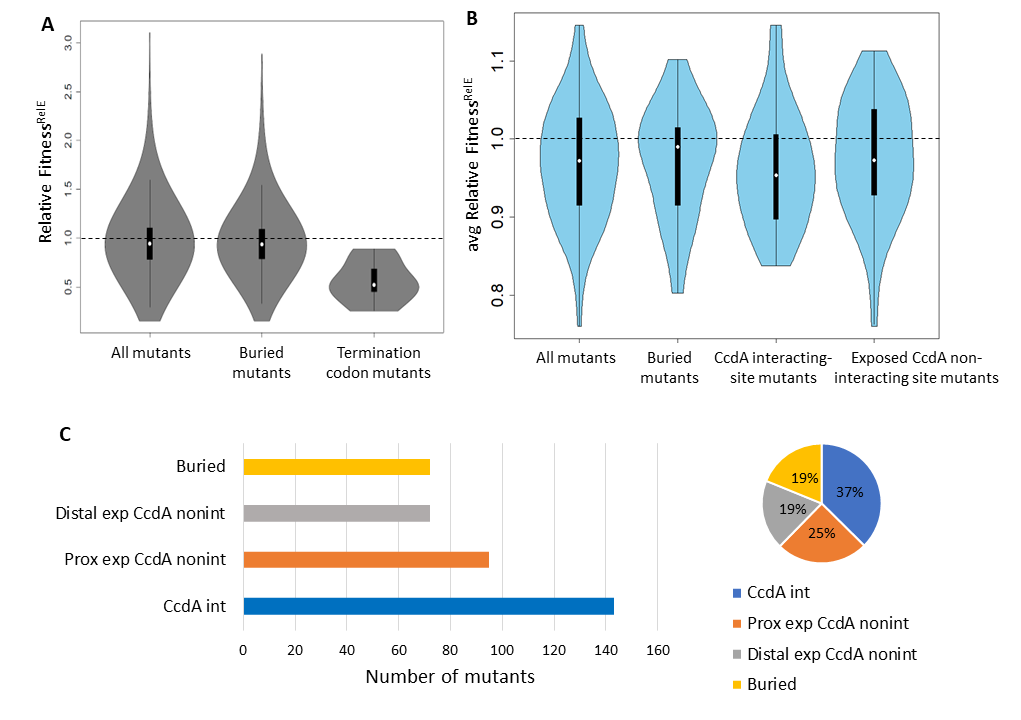

**Figure S2: The majority of CcdB mutants in the protein core show an active phenotype when expressed in an operonic context.** (A) Frequency distribution of RF^RelE^ for the entire dataset, for buried site mutants and TAA termination codon mutants. The majority of the buried-site mutants show a phenotype similar to the WT while the highest frequency stop codon in the *E.coli* genome, TAA gave the expected derepressing phenotype. (B) Distribution of RF^RelE^ scores averaged over each position for four structural categories, namely all mutants, buried site mutants, CcdA interacting site mutants, and exposed CcdA noninteracting site mutants. In (A-B), the width of the violin plot is proportional to the relative number of mutants. The white dot in the middle represents the median of the distribution. The box plot in black colour represents the inter-quartile range. The black dashed line corresponds to the normalised score obtained by the WT (RF^RelE^ = 1). (C) CcdA binding defective mutants with RF^RelE^<0.7, classified into four different structural classes, i.e., at the CcdA binding site (CcdA int), exposed and proximal (within 8Å) to the CcdA binding site (Prox exp CcdA nonint), exposed but distal (>8Å) to the CcdA binding site (Distal exp CcdA nonint) and in the protein interior (Buried). The length of the bar is proportional to the total number of mutants for each class (left panel). These absolute numbers are converted into percent fractions shown in a pie chart (right panel)

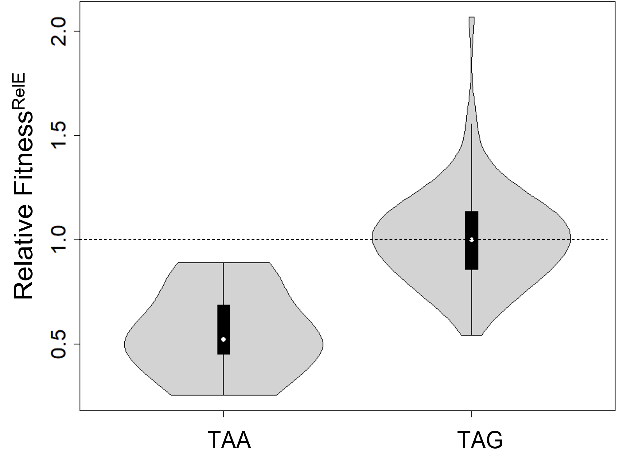

**Figure S3:** **Frequency distributions of RF^RelE^ for two different types of nonsense mutants (TAA and TAG).** The high frequency TAA stop codon mutant (also shown in Figure S2A) gave a derepressing phenotype whereas the TAG stop codon mutant spanned the entire range of scores for reasons not clearly understood.

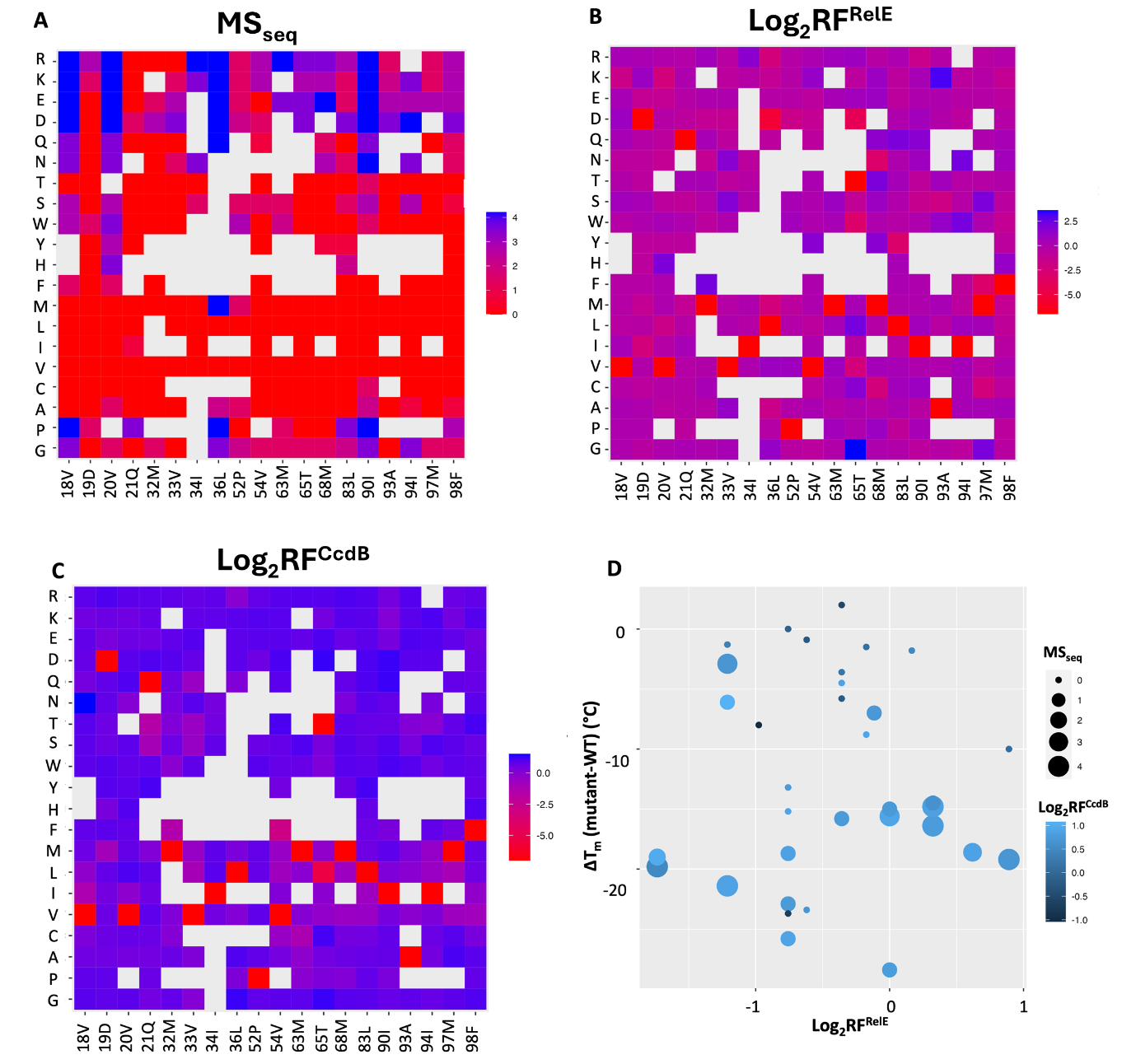

**Figure S4: Mutational effects on CcdB activity inferred from phenotypic screening and deep sequencing, of CcdB buried-site mutants in the absence and presence of CcdA.** MS_seq_, RF^RelE^ and RF^CcdB^ scores obtained from deep sequencing are averaged over synonymous mutants. For comparison, mutational scores were converted to z score and further represented relative to the WT z-score value. (A-C) On the left vertical axis, residues are grouped into (G, P), aliphatic (A–M), aromatic (F–W), polar (S–Q) and charged (D–R) amino acids. Residue numbers with their corresponding WT amino acid and substitutions are indicated on the horizontal and left vertical axes, respectively. Mutations where there is no data are shown in light grey. The colour scheme for the heat map is shown on the right of each plot, and the colour bar represents the relative z-scores. Relative z score for (A) MS_seq_ values for buried site mutants. Gradation of red to blue colour represents increasing relative z scores and decreasing CcdB activity. WT residue at each position is indicated in red, (B) RF^RelE^ scores for buried site mutants. Red to blue gradation represents increasing relative z scores, indicating increasing amounts of CcdAB complex. The WT residue at each position is indicated in red, (C) RF^CcdB^ for buried site mutants. A relative z score could either be because CcdB is neutralised by CcdA or it remains misfolded even in the operonic context. Gradation of red to blue corresponds to increasing relative z score. WT residue at each position is indicated in red colour. (D) Thermal stabilities of characterised buried-site mutants (3) were compared with their corresponding relative RF^RelE^, MS_seq_ and RF^CcdB^ scores. Coordinates (x,y) of each point reflect the corresponding relative RF^RelE^ value and the difference in thermal stability of the mutant with respect to WT (ΔT_m_). The size of the bubble increases with increasing MS_seq_ score. The blue colour becomes lighter with increasing RF^CcdB^ score. Smaller dots (lower MS_seq_ score) are typically darker (lower RF^CcdB^ score). Several highly destabilized mutants (ΔT_m_<-10^o^C) have RF^RelE^>1, indicating that they are well folded when expressed in the operonic context.

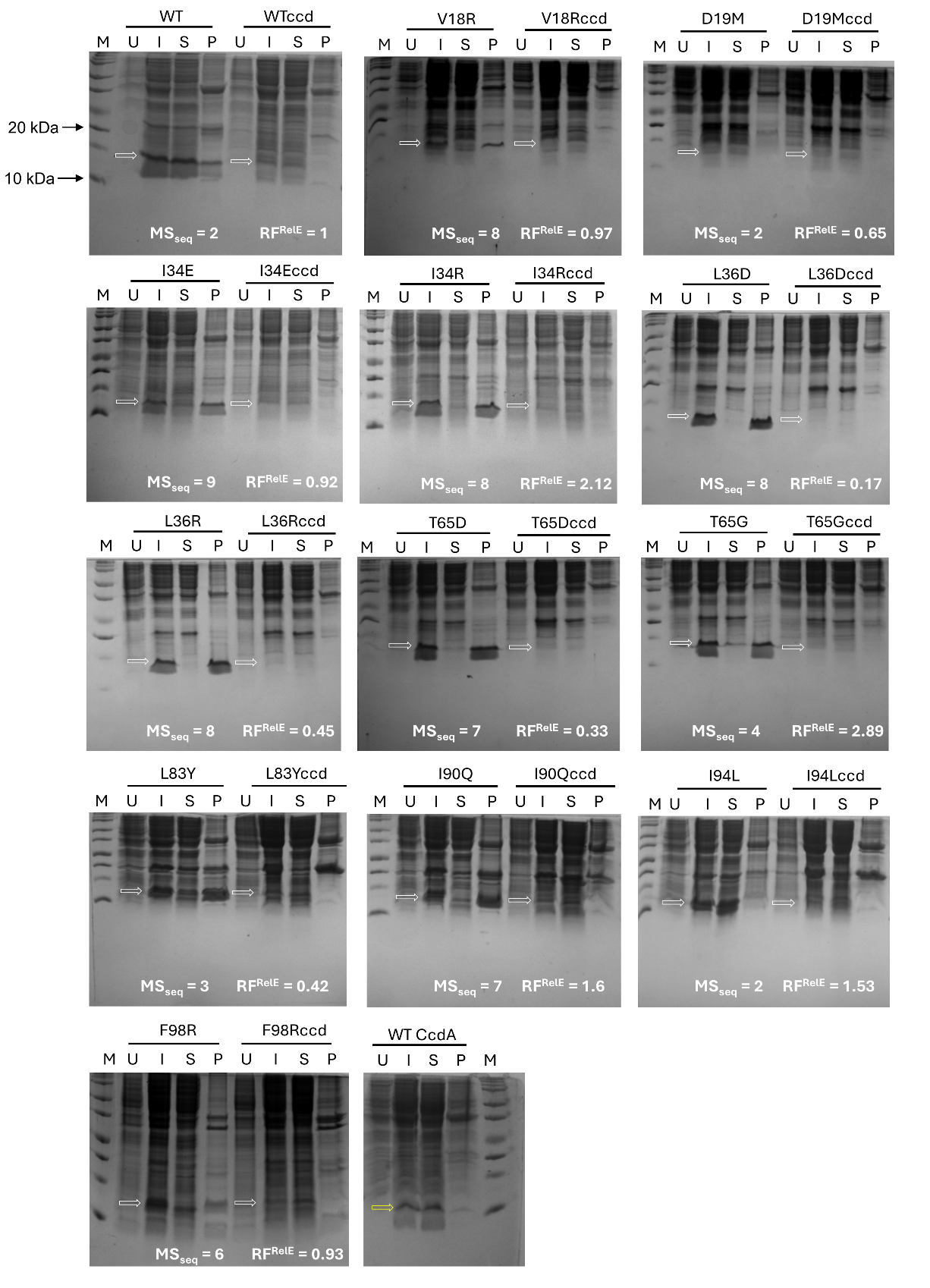

**Figure S5: Expression of buried-site CcdB mutants in the absence and presence of CcdA.** *In vivo* solubility of destabilised CcdB mutants when expressed alone relative to when they are present in an operonic context from an arabinose inducible promoter. The corresponding mutational score obtained from deep sequencing, i.e., MS_seq_ (-CcdA) and RF^RelE^ (+CcdA), for each mutant is mentioned on the gel. M represents the protein ladder. U, I, S and P represent uninduced, induced, supernatant and pellet cell lysate fractions, respectively. The fraction of protein in supernatant or pellet was determined by densitometric analysis following 15% Tricine SDS-PAGE and Coomassie staining. The white arrow indicates the likely position of the protein of interest (CcdB). The yellow arrow indicates the position of WT CcdA protein when expressed separately from an IPTG inducible promoter (bottom last panel). This assay was carried out in biological duplicates.

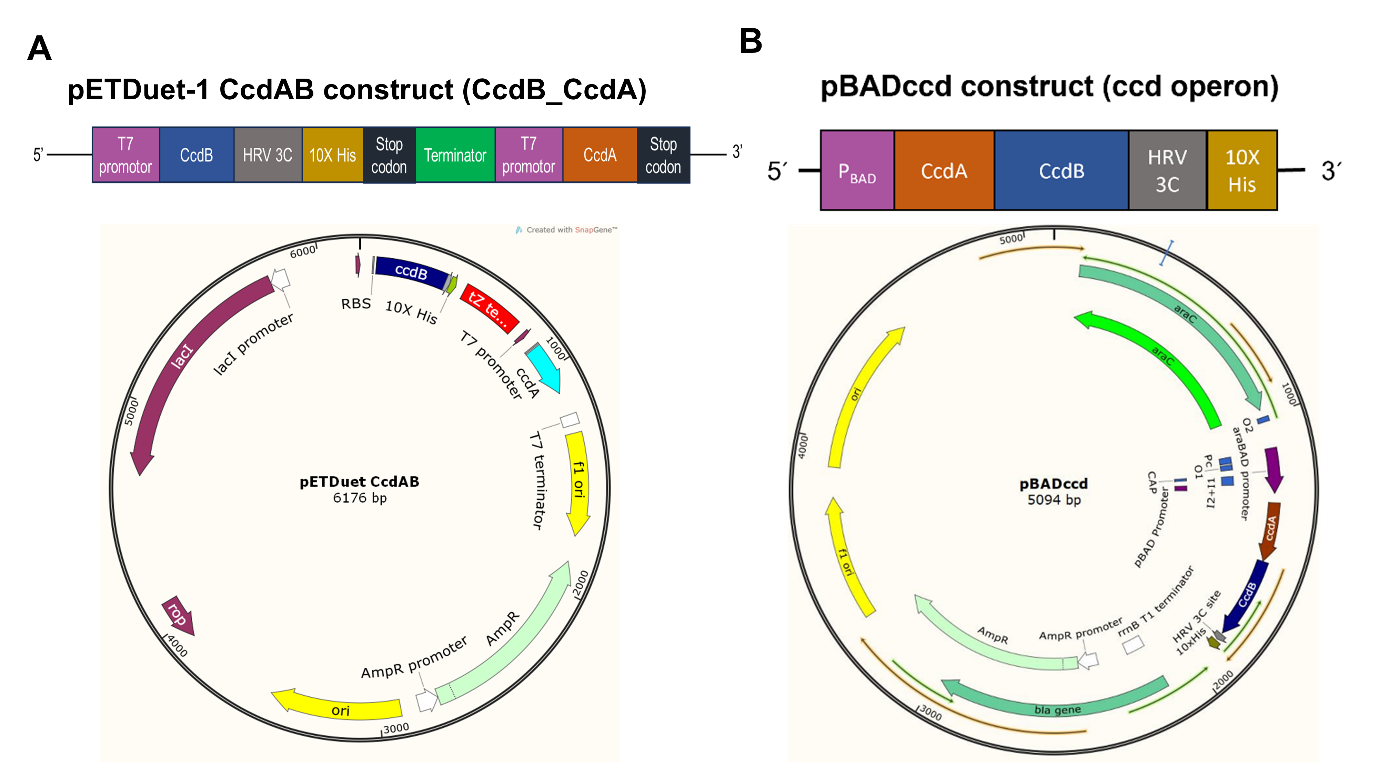

**Figure S6: Two different vector systems were used to perform *in vivo* solubility assays and for purifying proteins.** (A) pETDuet construct. Top panel shows *ccdB* and *ccdA* gene organization in the pETDuet vector. Here *ccdB* and *ccdA* are expressed separately under two different IPTG inducible T7 promoters in the same vector. The *ccdB* gene sequence is identical to that used in the pBADccd construct. Bottom panel shows the vector map of the construct. (B) pBADccd construct. Top panel shows gene organization of the *ccdAB* operon with a cleavable C-terminal His tag containing an HRV3C protease site in pBAD24 vector . ccdA and ccdB induction is under control of the arabinose inducible P_BAD_ promoter. Bottom panel shos vector map of the construct.

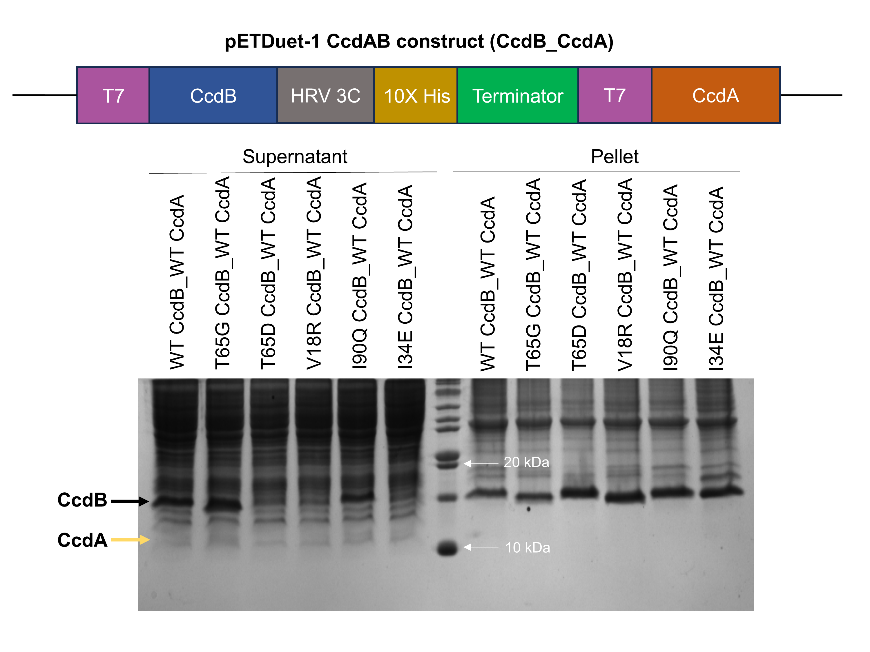

**Figure S7: Coomasie stained SDS-PAGE of His-tagged CcdB mutants in the presence of CcdA, expressed from different mRNAs.** *In vivo* solubility of Wild-type and destabilized CcdB mutants when expressed in a non-operonic context. Both CcdB and CcdA are expressed from two different T7 promoters. Black arrow indicates the CcdB protein and yellow arrow indicate CcdA protein. In order to visualize the CcdA bands, larger amounts of sample were loaded in this gel relative to those in Figure 2B.

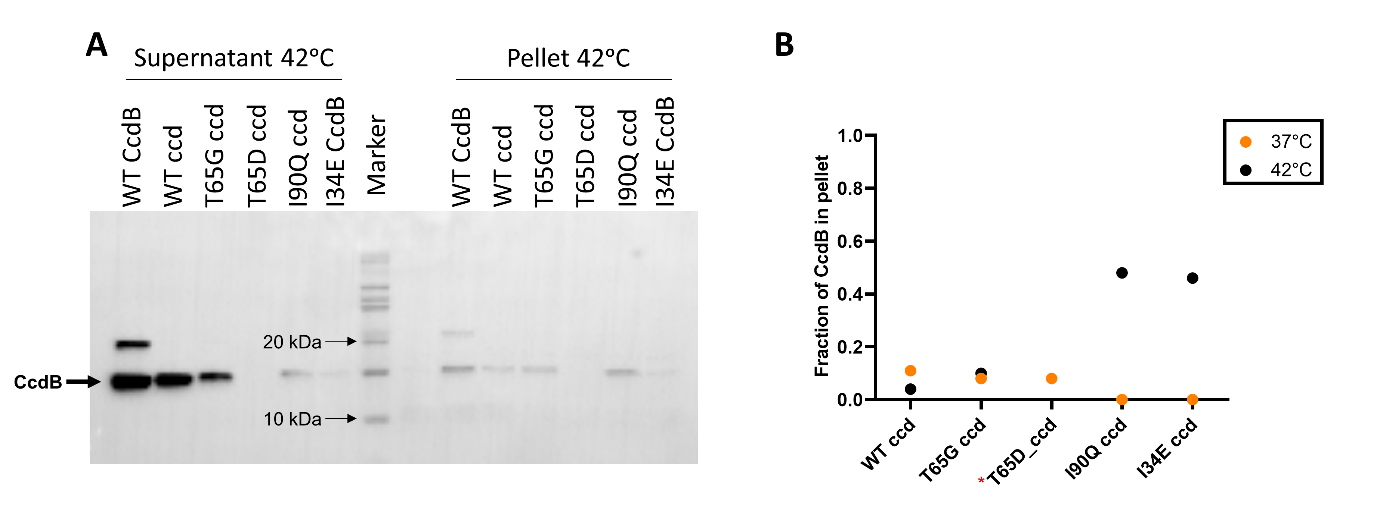

**Figure S8: Western blot of His-tagged CcdB mutants in the presence of CcdA.** *In vivo* solubility of Wild-type and destabilized CcdB mutants when expressed in an operonic context from an arabinose inducible promoter (A) at 42^ᵒ^C. (B) Comparison of fraction of CcdB in pellet relative to supernatant at 37^ᵒ^C and 42^ᵒ^C. The arrow indicates the protein of interest (CcdB). Data for 37^ᵒ^C are shown in Figure 2A. (* represents no detectable band in the operonic context at 42°C)

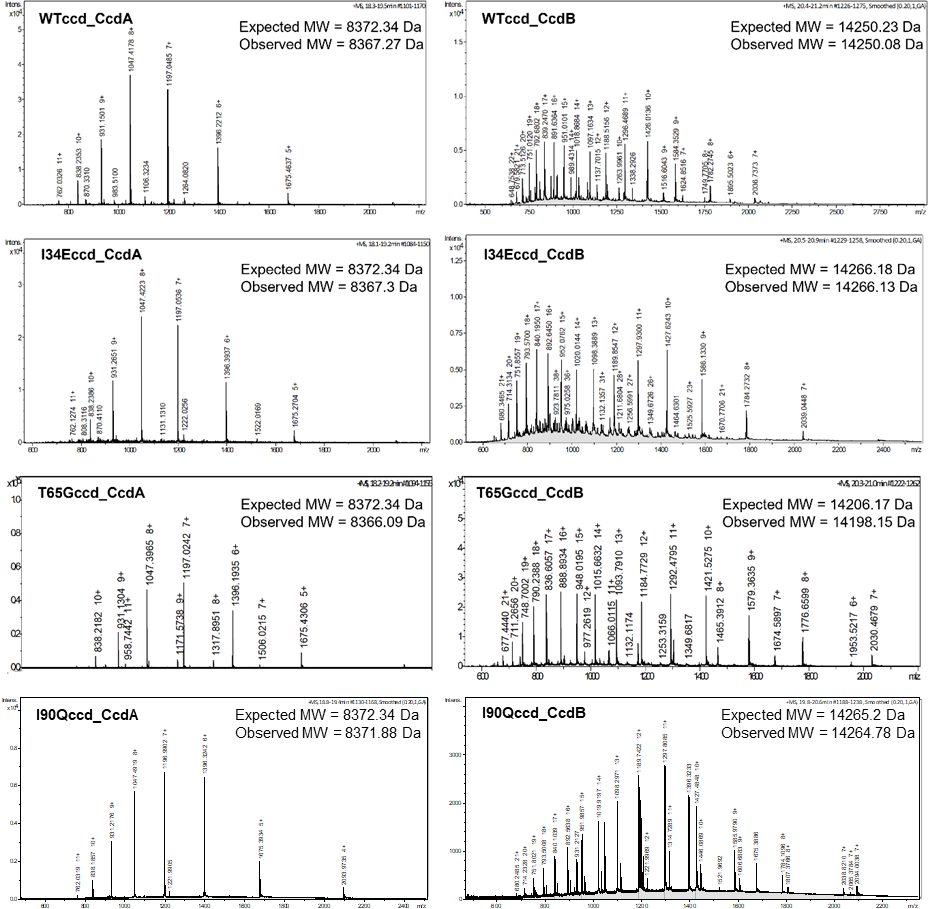

**Figure S9: Mass spectrometry analysis of purified WT and mutant complexes when CcdA and CcdB are expressed in an operonic context.** ESI Mass spectra of CcdA and CcdB protein components from mass spectrometry of CcdAB complex for WT and CcdB mutant (I34E, T65G and I90Q) proteins. Spectra for V18R could not be recorded because it could not be purified in sufficient amount due to its poor expression. In all cases the CcdB has a C-terminal His tag. Expected and observed Molecular Weights (MW) for the CcdB and CcdA components are mentioned above the spectra.
